# Dynamic Regulation of the Immune Repertoire of Bacteria

**DOI:** 10.1101/2025.09.17.676245

**Authors:** Zhi Zhang, Sidhartha Goyal

## Abstract

The CRISPR-Cas system provides adaptive immunity in many bacteria and archaea by storing short fragments of viral DNA, known as spacers, in dedicated genomic arrays. A longstanding question in CRISPR-virus coevolution is the optimal number of spacers for each bacterium to maintain proper phage coverage. In this study, we investigate the optimal CRISPR memory size by combining steady-state immune models with dynamical antigenic traveling wave theory to obtain both analytic and numerical results of coevolutionary dynamics. We focus on two experimentally supported phenomena that shape immune dynamics: primed acquisition, where partial spacer-protospacer matches boost acquisition rates, and cassette expansion, where a short-term increase in memory size drives population dynamics. We find that under primed acquisition, longer optimal arrays benefit from maintaining multiple, partially matching spacers. In contrast, dynamic cassette expansion favors shorter arrays by amplifying the fitness advantage of acquiring a few highly effective new spacers. Together, our results highlight that memory optimality is not fixed, but instead shaped by the interaction of acquisition dynamics and population-level immune pressures.

## I. INTRODUCTION

Coexistence and defense against rapidly adapting viruses is a fundamental challenge across all domains of life, from bacteria to humans. While generalized innate immune systems provide broad protection against all viruses, they are likely not as effective as mechanisms that specifically learn and target particular pathogens. While the adaptive immune systems of vertebrates learns through the generation and selection of antibodies with high binding affinity to antigens, many bacteria and archaea species evolved an adaptive immune strategy through the CRISPR-Cas system, which specifically targets viral DNA [1–3]. Over all the different types of CRISPR-Cas mechanisms, the general mechanism all rely on arrays of clustered regularly interspaced short palindromic repeats (CRISPR) loci. These loci consist of repetitive DNA sequences interspersed with short, unique sequences known as spacers, which are homologous to regions of previously encountered bacteriophage (phage) genomes. Immunity is achieved by transcribing each spacer into an RNA molecule (crRNA) to guide CRISPR-associated (Cas) proteins, which will cleave the recognized nucleic acids, thereby neutralizing the threat of foreign phage elements. The phage-derived DNA segments targeted by the CRISPR system before spacer integration are hereby referred to as protospacers.

The optimal number of spacers in each CRISPR locus remains a central question in the understanding of CRISPR-virus coevolution. Laboratory strains of bacteria [4, 5] often begin without spacers to acquire just a few spacers throughout the experiment. However, bacterial strains isolated from natural environments frequently have longer CRISPR cassettes, which may contain a few dozen to a few hundred spacers [5–7].

In this work, we assume that large-scale recombination occurs frequently enough within these populations to justify treating all spacers as part of a shared population pool. This regime may be particularly relevant in hot-spring microbial mats, where recombination contributes to genetic diversity on a scale comparable to mutation [8, 9] while the highly conserved spacer profiles suggest their CRISPR immune cycle operates on the order of years [10]. Our focus is on the long-term dynamics of CRISPR memory size in coevolving phage-bacteria systems under conditions of high horizontal gene transfer (HGT) rates, which are known to occur in hot-springs environments [11]. Furthermore, we assume a single CRISPR-Cas type across all individuals in the environment.

In such environments, prolonged coexistence between phage and host populations facilitates the advent of a coevolutionary steady-state, while HGT introduces variability in the effective memory size. Understanding how these forces interact to shape the optimality of CRISPR loci offers new insight into the diversity of immune strategies observed in natural microbial communities.

The optimal number of spacers in each CRISPR loci remains a central question in the understanding of CRISPR-virus coevolution. Laboratory strains of bacteria [4, 5] often begins without spacers to acquire just a few spacers throughout the experiment. However, bacteria strains isolated from natural environments frequently have longer CRISPR cassettes which may contain a few dozen to a few hundred spacers [5–7].

In this work, we assume that large-scale recombination occurs frequently enough within these populations to justify treating all spacers as part of a shared population pool. This regime may be particularly relevant in hot-spring microbial mats, where recombination contributes to genetic diversity on a scale comparable to mutation [8, 9] while the highly conserved spacer profiles suggest their CRISPR immune cycle operates on the order of years [10]. Our focus is on the long-term dynamics of CRISPR memory size in coevolving phage-bacteria systems under conditions of high horizontal gene transfer (HGT) rates which are known to occur in hot-springs environments [11].

In such environments, prolonged coexistence between phage and host populations facilitates the advent of a coevolutionary steady-state, while HGT introduces variability in the effective memory size. Understanding how these forces interact to shape the optimality of CRISPR loci offers new insight into the diversity of immune strategies observed in natural microbial communities.

Previous models have shown that increasing the size of the CRISPR-Cas memory can lead to diminishing immune effectiveness due to intrinsic constraints on the number of interference complexes and the sensitivity of the CRISPR machinery [12]. Moreover, overexpression of CRISPR-Cas components has been associated with toxic side effects, particularly due to the risk of autoimmunity through self-targeting [13]. These arguments do not help to understand important phenomena in which cassette sizes are dynamic and change substantially within bacterial populations experiencing sustained selective pressure from rapidly evolving phage. In this work, we focus on two mechanisms that can drive such dynamics: primed acquisition and cassette size expansion mediated by temporal regulation of spacer acquisition and loss rates.

**Primed Acquisition** refers to a phenomenon where an already acquired primed spacer biases the acquisition probability of future protospacers, which generally leads to substantially increased acquisition rate [14, 15]. This effect has been observed across CRISPR subtypes and bacterial strains [16–19]. Accurately modeling primed acquisition has posed significant challenges, as it requires simulating bacterial populations carrying multiple spacers per individual alongside a highly diverse distribution of phage protospacers, while also incorporating acquisition biases informed by sequence similarity.

**Cassette Expansion** is a result of the mechanisms of acquisition and loss of spacers. Even from the earliest experiments, it has been shown that CRISPR cassettes can rapidly expand their number of spacers within a few generations when under phage threat [20]. However, regulation of CRISPR-Cas systems is essential since these systems are both metabolically expensive and pose a risk of autoimmunity [21]. In our model, we incorporate a tradeoff whereby temporary increases in acquisition efficiency are allowed at the cost of decreased net future spacer acquisition. Unlike previous models, which assume a fixed or averaged memory size across the population, we explore how time-dependent variation around optimal memory size affects both bacterial and phage populations in a coevolutionary setting.

Multiple experiments have shown that CRISPR interference can still occur despite mismatches of a few base pairs between spacer and protospacer sequences [22–24]. This immune cross-reactivity have previously been modeled using agent-based simulations that explicitly track the evolution ofindividual spacer and protospacer sequences [25]. Similar agent-based models have been used to successfully model both theoretical and experimental observations of CRISPR-Cas systems [26]. However, rather than relying solely on agent-based simulations, we first show that in the regime of high cross-reactivity, these simulation results can be embedded into a low-dimensional space that preserves spacer–protospacer distance metrics. Such a representation allows us to construct a predictive model of coevolution. We then formulate a partial differential equation (PDE) model onto this representation that yields analytical insight into the two phenomena described above.

More specifically, we combine two complementary theoretical approaches to investigate optimal CRISPR memory size. First, we borrow from the statistical view of CRISPR–virus interactions, used to compute an optimal number of spacers under simplified biochemical assumptions. Second, we used a time-dependent dynamical framework based on traveling wave theory to capture the coevolutionary immune dynamics. We demonstrate that primed acquisition tends to favor larger memory sizes, as individual spacers become less effective at interference, thereby requiring greater redundancy. In contrast, cassette expansion dynamics tend to favor smaller memory sizes, as bacteria with fewer spacers can acquire new, more effective spacers, which can rapidly drive out phages in the environment.

## II. MODEL AND METHODS

### A. Antigenic Embedding of Spacers

Starting from a theoretical paper [27], agent-based simulations shows that a stable coevolutionary regime can can emerge in which neither population completely outcompetes the other. This regime is specifically modulated by the cross-reactive immunity, which maintains a delicate drifting equilibrium where both populations evolve in lockstep. Although phage protospacers mutate randomly in an approximately isotropic manner, selective pressure from bacterial spacers drives protospacer evolution toward sequences with lower immune coverage. The number of such escape sequences is greatly reduced with increased cross-reactivity. As a result, a cyclical arms race emerges in which spacer adaptation promotes protospacer escape, which in turn selects for new bacterial spacers.

We show that agent-based simulation data with cross-reactivity can be represented in at most two dimensions while preserving the relevant distance structure. Starting from the sequence information generated by the simulations, we applied metric multi-dimensional scaling (MDS) to embed the Hamming distance matrix of all unique spacers and protospacers (see Fig. 1A). Metric MDS was chosen because it aims to preserve the metric relationships of the original sequence space (Hamming distance) within the embedded Euclidean space.

**FIG. 1.**
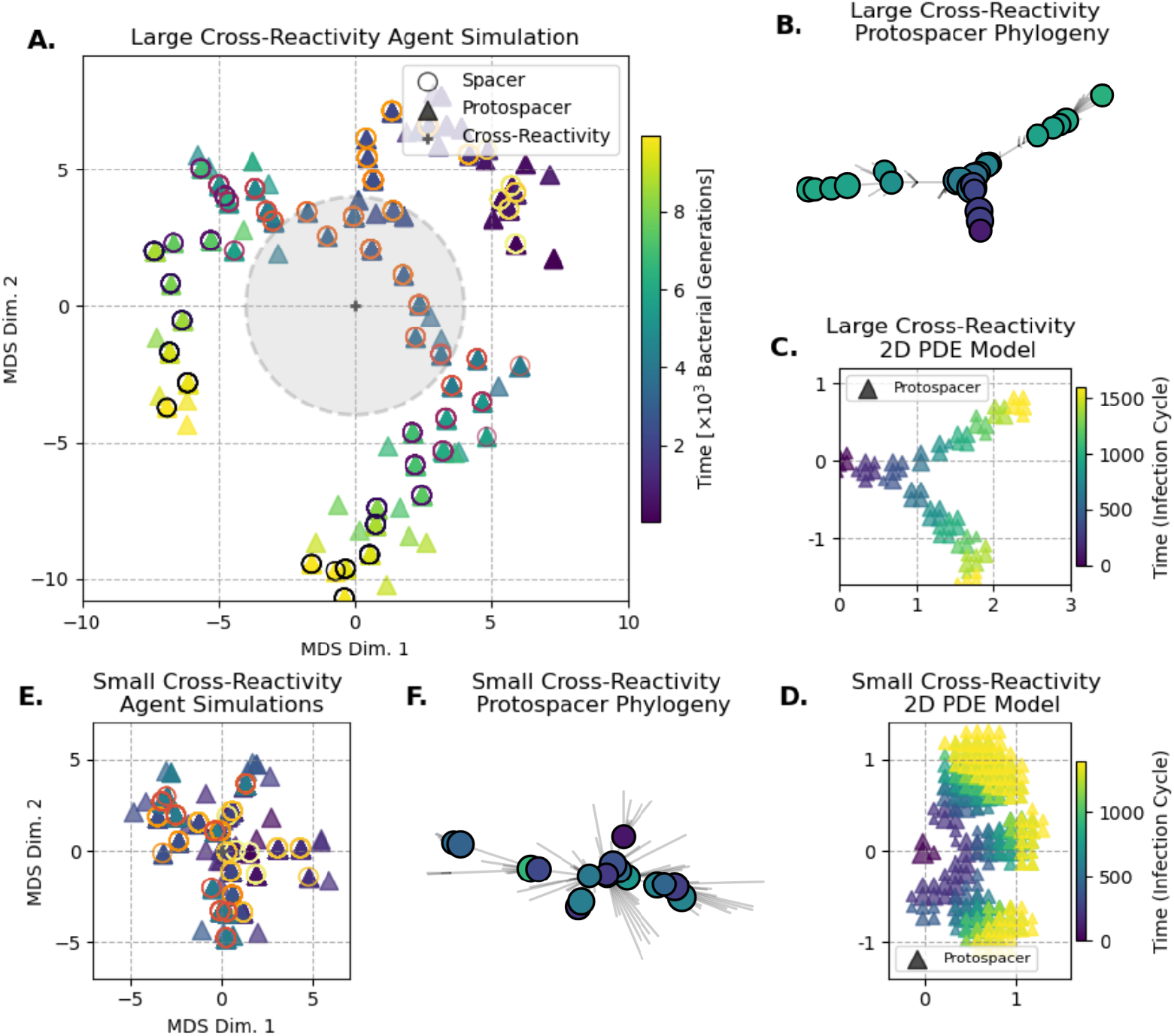
Comparison of Large vs Small Cross Reactivity Simulation in Agent-Based vs PDE Models (A) MDS embedding of agent-based simulations, the relevant comparable parameters are exponential cross-reactivity radius *r* = 4 and Poisson mutation rate *µ* = 10^*−*5^. This simulation clearly bifurcates into two lineages, which are visible only in 2D and above. Distances are roughly the same to the number of base pair substitutions. (B) Unrooted Phylogenyc tree of the most abundant protospacer sequences generated during the large cross-reactivity agent-based simulations. Note that the color of each node uses the same colorbar as fig. A and a clear chronological order can be seen along each of the two lineage branch (C) PDE model with large cross-reactivity, *r* = 1bp and *µ* = 10^*−*5^, similar to the agent-based model, this simulation splits into two distinct lineages. (D) PDE model with small cross-reactivity *r* = 0.1, no distinct lineages, protospacers diversify almost immediately to escape spacer immunity. (E-F) MDS embedding and phylogenetic tree for agent-based simulation with no cross-reactivity (single mutation escape). Note how simulation doesn’t have a specific evolution direction in the embedded space, the phylogeny also does not have any chronological ordering.

Ideally, a distance of one between two embedded spacers corresponds to a single nucleotide substitution in the original 30 bp sequences. We also attempted to apply the same embedding procedure to several experimental datasets [4, 28, 29]. However, these attempts were inconclusive due to limited temporal resolution and insufficient sequencing depth of those experiments. A more detailed discussion of the available experimental evidence, along with embedding results for one dataset [28], is provided in the Supplementary Information (see SI Sec. 3B).

Examining the embedded simulation results, we observe that individual lineages predominantly move along a single dimension (see Fig. 1A). However, a purely one-dimensional embedding of all sequences becomes inaccurate when multiple lineages coexist simultaneously (see SI Sec. 3A). A detailed analysis of embedding errors and drift prediction is provided in the Supplementary Information, but in short summary: Increasing the number of embedding dimensions reduces the total model error. However, the most predictive models are made with smaller dimensional embeddings, especially when the dataset is decomposed into individual lineages. Importantly, these 1-dimensional lineages can only be clearly separated when the embedding has at least two dimensions. Taken together, these observations suggest that two dimensions provide the optimal balance between representational accuracy and predictive power.

An useful indicator of whether a low-dimensional embedding is appropriate is the phylogenetic structure of spacer or protospacer sequences constructed from sequence distance metrics. At large cross-reactivity, the phylogenetic tree of protospacer evolution is quite simple and forms long chains of single branches, where each new sequence appears chronologically according to its establishment time (see Fig. 1B). In effect, cross-reactivity limits the number of branching lineages that can emerge at each generation (see Fig. 1E). By contrast, a small (or binary) cross-reactivity radius promotes protospacers diversification at each generation and thus making a low-dimensional embedding impossible. In this small cross-reactivity regime, the phylogeny shows rapid diversification away from the ancestral strain, with no clear chronological ordering within branches (see Fig. 1F) and thus indicating that no such embedding is possible.

### B. Adaptive Immunity Coevolution

In the previous section, we discussed how results from agent-based simulations could be represented in a low-dimensional spacer. In this section, we propose a partial differential equation (PDE) framework that captures the large-scale behavior of bacterial and phage populations by simplifying their interactions onto a shared low-dimensional embedding space. While individual-level descriptions are lost, this model offers a powerful means to investigate the impact of larger memory size and other population-scale parameters while providing an independent population simulation scheme. Similar PDE-based models were originally developed to describe viral evolution driven by shifting immunity [30–32].

We begin by assigning position vectors in a projected Euclidean space to each unique spacer–protospacer sequence pair. This theoretical embedding space will hereafter be referred to as the antigenic space, following the terminology of [31]. In practice, such an antigenic space can be constructed using low-dimensional embedding techniques similar to those described in the previous section. Within this space, we represent the protospacer population by a density field *n*(*x, t*) where *x* denotes the position vector in the embedded space associated with a matching spacer/protospacer sequence. The evolution of the protospacer population can then be described by a continuous mean-field equation that captures the effects of isotropic mutation and fitness-driven selection. In this framework, each infection cycle is assumed to occur on a timescale of roughly one bacterial generation (Fig. 2A.).

**FIG. 2.**
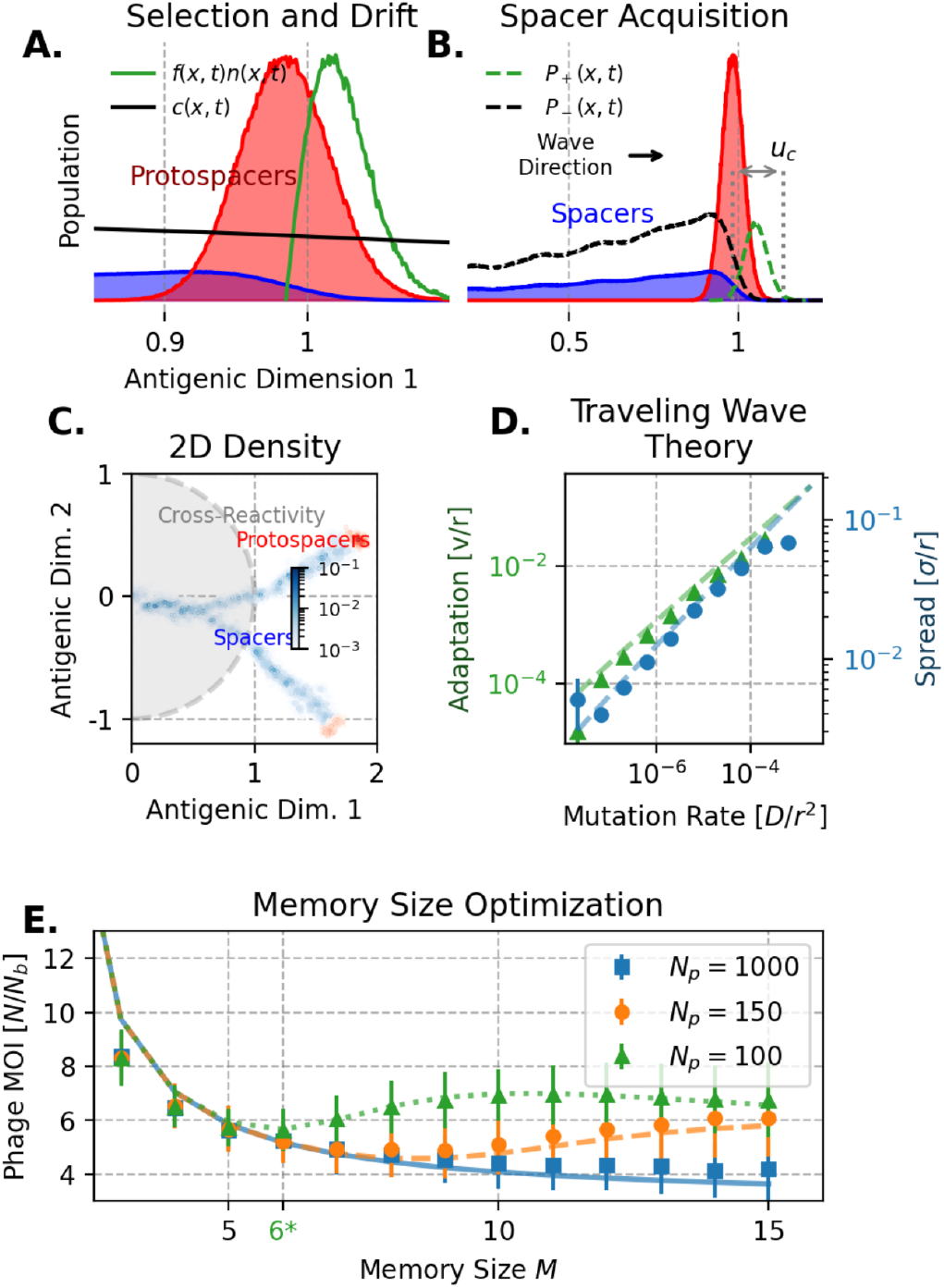
Model for phage-bacteria coevolution captures biochemical limitation in a dynamically stable equilibrium. Simulations were done with *r* = 1, *R*_0_ = 100 and *α* = 0.01. (A) Schematic of traveling linear fitness waves undergoing selection and drift. The linear fitness regime is called as such because the coverage *c*(*x, t*) can easily be linearized in this regime. (B) A more zoomed out view of spacer wave, note how the spacer population covers more antigenic distance compared to the protospacer population. Note how the acquisition probabilities also correspond to the current population distribution. (C) Population density view of traveling waves in 2 dimensions (same simulation as fig.1 C using *D/r*^2^ ≈ 10^*−*5^). Note the long tail of spacers following the current protospacer populationn (C) Coexistence conditions for diversity and adaptation speed. Only around those specific population spread and velocity can a traveling wave exist. (D) Dependence on M/Cas proteins on the amount of interference protein and binding sensitivity, there exists an optimal amount of spacers per bacterium that minimizes the coevolving phage population. Note that a linear correction was needed to correctly match simulation to theory, where *N*_sim_ ≈ 0.8 · *N*_theory_ .

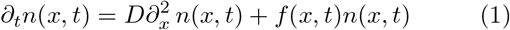

After each infection cycle, individual phages in the population mutate by randomly jumping to a new location in antigenic space. At the population level, this process generates an effective diffusion term with diffusivity *D*. By using the cross-reactivity radius to define a nucleotide-length scale, the normalized quantity *D/r*^2^ can be used as a protospacer mutation rate. Phage mutation rates per base pair, which have been estimated to be from 10^−8^ to 10^−6^ substitutions per nucleotide per infection cycle (s/n/c) for DNA phages and from 10^−5^ to 10^−3^ for RNA phages [33, 34]. These higher RNA mutation rates are relevant because some CRISPR subtypes are capable of targeting RNA phages [35]. In this study, we will be working with mutation rates *D/r*^2^ in the range of 10^−6^ to 10^−5^.

Our smallest PDE simulation length scale was 10^−3^bp. We argue that the fine-grained approximation of mutational effects in Eq. 1 remains valid, even though single substitutions in the protospacer sequence correspond to relatively large jumps in the antigenic space. In addition, snapshot distributions from the agent-based simulations show that spacer and protospacer populations remain highly localized (see SI Sec. 3A). Furthermore, the fitness distribution of bacterial populations is expected to be much more fine-grained than their underlying genetic distribution due to the influence of many non-genetic factors [36]. As such, we intentionally use a resolution in antigenic space that is finer than the genetic scale in order to accurately capture the evolution of the population distribution over generational timescales.

The bacteria population, or more specifically the spacer population density, is denoted by *n*_*s*_(*x, t*). In our model, the bacteria population *N*_*b*_ is constant and not subject to phage selection. Events that disrupt coexistence, such as population runoff/extinction, will be described entirely by the change in the total phage population size *N* (*t*) = ∫ *n*(*x, t*)*dx*. The total spacer population size is given by *N*_*b*_*M* = ∫ *n*_*s*_(*x, t*)*dx* where *M* is the average number of spacers per bacteria. We fixed the expression level of the CRISPR system and assumed that all regulations that occur are averaged on a population scale. The evolution of the spacer population is governed by the acquisition and loss of spacers, which occur at the same infection cycle time.

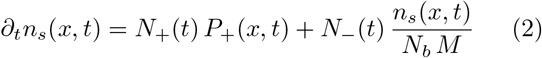

Here, the acquisition rates *N*_+_(*t*) and *N*_−_(*t*) are simply the total number of spacers acquired and lost at each cycle. These rates are proportional to the current phage population *N* (*t*). The base case, without net memory gain or loss, is assumed to be *αN* (*t*) where *α* sets the per-cycle acquisition rate of protospacers. Throughout our simulations, this rate is usually set to *α* = 0.01, which corresponds to the rate at which 1% of the total protospacer population will be converted to spacers at each timestep, *eg*. at MOI=10 and *α* = 0.01, 10% of all bacteria will acquire a new spacer at each generation. While CRISPR acquisition rate varies widely depending on the species, subtypes, and environment [37, 38], the choice of *α* = 0.01 is motivated by typical phage-to-bacteria ratios (or multiplicity of infections, MOI) observed in natural systems [39] and coevolution experiments [**?** ]. *P*_+_(*x, t*) is the acquisition probabilities each unique spacer. The unbiased acquisition probability is directly proportional to the phage density: *P*_+_(*x, t*) = *n*(*x, t*)*/N* (*t*). On the other hand, spacer loss is proportional to spacer abundance and we assume all spacers have an equal rate to be lost. Furthermore, we do not assume a preferential ordering of spacers within a cassette; the older spacers will naturally fall off from the immune population as the two populations move within the antigenic space through each infection cycle.

Finally, the last key element to consider is the collective coverage *c*(*x, t*) of the spacer population, which is the total effect of all spacers on targeting phages across the antigenic space. This coverage function accounts for cross-reactivity. To model the decrease in CRISPR effectiveness with mutational distance, we consider an exponential cross-reactivity kernel with radius *r*.

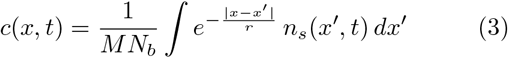

Based on previous work done with similar models [31, 40], large cross-reactivity: *r* ≫ *δx* in this PDE model will also lead to to a canalized evolutionary trajectory between host and pathogens (see Fig. 1C). In contrast, when the cross-reactivity radius is small: *r* ≪ *δx*, the dynamics instead produce radial expansion away from the origin (see Fig. 1D). Experimental measurements of CRISPR cross-reactivity suggest that a reasonable choice is *r* = 1000*δx* = 1, which corresponds to a single point substitution difference in the spacer genetic sequence (Fig. 2C).

### C. Fitness Model

We choose the fitness function of eq. 1 for the system that incorporates the fundamental constraints of the bacterial biochemical machinery. Previously proposed in 12 for phage and bacteria population in equilibrium, we model the probability of successful infection of a single bacterium (Eq. 4) as a function of the expected number of effective spacers in its CRISPR cassette (accounting for cross-reactivity). The expected number of effective spacers is the previously mentioned coverage *c*(*x, t*) of the spacer population. In our model, the infection probability decreases with the number of effective spacers (including the effects of cross-reactivity) present in each bacterium at infection. However, even in the presence of a single effective spacer, a successful immune response is not guaranteed, as it will depend on the probability of expression and interference. For our purposes, we only consider the case where infection can occur only when one or fewer effective spacers are present. While a higher number of effective spacers can be considered, their contribution to the infection probability is very small in the relevant regime (SI Sec. 1C).

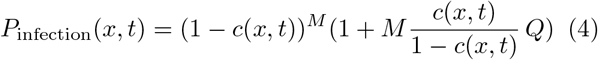

Here, *Q* denotes the probability of unsuccessful interference despite the presence of a single effective spacer in the CRISPR cassette. *Q* is uniquely determined by the expression level of our CRISPR arrays, which depend on *N*_*p*_ which is the number of spacers expressed as Cas-complexes. A more detailed explanation of *Q* can be found in SI. Section TBD.

This non-linear infection probability effectively captures the tradeoff between memory size *M* and expression levels of *N*_*p*_. There exists an optimal memory size *M*, that scales along *N*_*p*_, which minimizes *P*_inf_ . Memory sizes below this optimum do not generate enough coverage against phages, while higher memory sizes dilute the expression of each spacer, effectively reducing interference. Although the exact value of this optimality does depend on the specific coverage *c*(*x, t*) from Eq. 4, this nonlinear effect was preserved in our dynamical simulations (Fig. 2E).

Finally, we adopt a simple fitness model in which the phage growth rate depends directly on the probability of successful infection. The key infection parameter is *R*_0_, which is the basic reproduction number of the phage. As shown in various experiments [41], this value seems to be approximately 100 for various species of lytic phage. While more detailed models of fitness, such as SIR-type frameworks, have been used for both bacteria-phage and antigenic evolution [40, 42], we opt for a more coarse-grained description of each infection cycle. Our focus is on the long-term coevolution of the system, allowing us to neglect the short-term fluctuations of subpopulations.

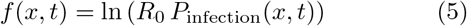

### D. Antigenic Traveling Waves

Solving the general system of Eq. 1 and Eq. 2 analytically is typically impossible, but there exist some initial conditions for which the general long-term behavior of the population can be inferred. One such non-trivial case is that of traveling waves, where the total proto-spacer population *N* remains constant and the population density takes the form *n*(*x, t*) = *n*(*x* − *vt*), corresponding to a co-moving frame with a steady velocity *v*. While many regimes of traveling waves exist, including the classical FKPP wave [43, 44]. These regimes have vastly different behaviors when extended to higher dimensional spaces. As shown earlier, agent-based simulations at large cross-reactivity can be embedded in a two-dimensional space, with individual evolutionary lineages effectively constrained to one dimension. In this work, we therefore focus on a specific traveling-wave regime: the linear fitness regime [31]. This regime has been previously shown to produce similar long, canalized evolutionary trajectories in two dimensions (Fig. 2B).

As previously discussed, the linear fitness regime typically emerges when the cross-reactivity radius is much larger than the mutation step size: *δx* ≪ *r* [40]. In this limit, the fitness landscape experienced by the phage population can be locally approximated as linear. An additional advantage of this regime is that it allows analytical approximations for key quantities such as the phage adaptation speed *v* and phage coevolution cluster spread *σ*^2^ (Fig. 2D) [30]. Finally, the adaptation speeds predicted by our model, given the mutation rates considered here, fall within the upper observed range of 10^−4^ to 10^−3^ s/n/c reported for some phages [45].

Finally, this regime imposes two important constraints on the average coverage *c*^∗^ of the protospacer population. First, to ensure a consistent balance between spacer formation and phage escape and maintaining the longterm coevolution, the value of *c*^∗^ must be consistent with the wave adaptation speed and spacer turnover time, *τ* = *N*_*b*_*M/*(*αN* ). Second, since the protospacer population must remain constant, the net fitness of the average phage must be 0: *f* (*c*^∗^) = 0. This requirement implies that *c*^∗^ must also be the root of the fitness function defined in Eq. 5. Together, these two conditions constrain the system to a unique steady-state solution with a constant phage population *N* and also a constant multiplicity of infection (MOI) *N/N*_*b*_. A more detailed discussion of these constraints can be found in the Supplementary (see SI Sec. 1A).

To close our traveling wave framework, we need to validate some final assumptions of the linear fitness regime. First, *vδt* ≪ *r* is maintained since adaptation at each infection cycle typically involves sub-nucleotide scale shifts, below the cross-reactivity threshold. Second, the memory of spacers extends beyond the current spread in the protospacer population or *σ* ≪ *vτ* . We believe it to be a reasonable assumption given data that shows long spacer persistence time (up to years) in bacteria with large CRISPR cassettes. Finally, we define a key length-scale of *u*_*c*_ which represents the antigenic distance from the wave center to the most fit individuals, as those outliers play an important role in driving the adaptation [46].

Finally, the main motivation for focusing on the linear fitness regime is that it provides an effectively one-dimensional description of population dynamics, even when the underlying embedding space is higher dimensional. In this regime, the canalized traveling-wave trajectory remains localized along a narrow evolutionary path, including in the two-dimensional embedding observed in the agent-based simulations. This observation further supports the idea that individual evolutionary lineages can be effectively tracked along a single dimension. In this work, most of our theoretical analysis and simulations are performed in one dimension. These one-dimensional models should therefore be interpreted as describing the dynamics of individual evolutionary lineages embedded within a higher-dimensional antigenic space. We also verified that key results can be reproduced using two-dimensional simulations (see SI Sec. 3A), and simulation code is available for both dimensionalities.

## III. RESULTS

### Primed Acquisition

Primed acquisition is the phenomenon where new spacer acquisition is biased towards spacers that are genetically related to those already present in the cassette [47]. Priming can occur when a previously acquired spacer, even when it doesn’t apparently confer any immune benefit, can serve to guide the CRISPR machinery to preferentially acquire spacers towards either a protospacer that’s genetically similar or identical [17]. Although the precise mechanism of acquisition is still only partially understood, many experiments have been done for different CRISPR subsystems in different environments [16–19, 48]. In natural populations, we often see spacer pairs with highly similar genetic makeup following one another in an array ( ∼ 2% of arrays in E.Coli have repeated spacers according to [49]). Primed acquisition can also target protospacers located further on the phage genome. An oversight of this model would be that acquired protospacer could be genetically dissimilar while being acquired through a primed mechanism. However, looking at the acquired protospacer mapping from several experiments [17, 18, 50], they suggest that priming does not appear to target any new genomic regions beyond those observed in naïve acquisition. Priming overall results in a narrower distribution in acquisition sites compared to naive acquisition.

Our mathematical model for primed acquisition first is to change the protospacer acquisition probability *P*_+_(*x, t*) in a way that will result in a bias towards the current immune (spacer) population. This is modeled in a similar way to the cross-reactivity kernel, where each spacer in the immune population will bias the phenotypical acquisition profile *π*_+_(*x, t*) by its own distribution (Fig. 3A):

**FIG. 3.**
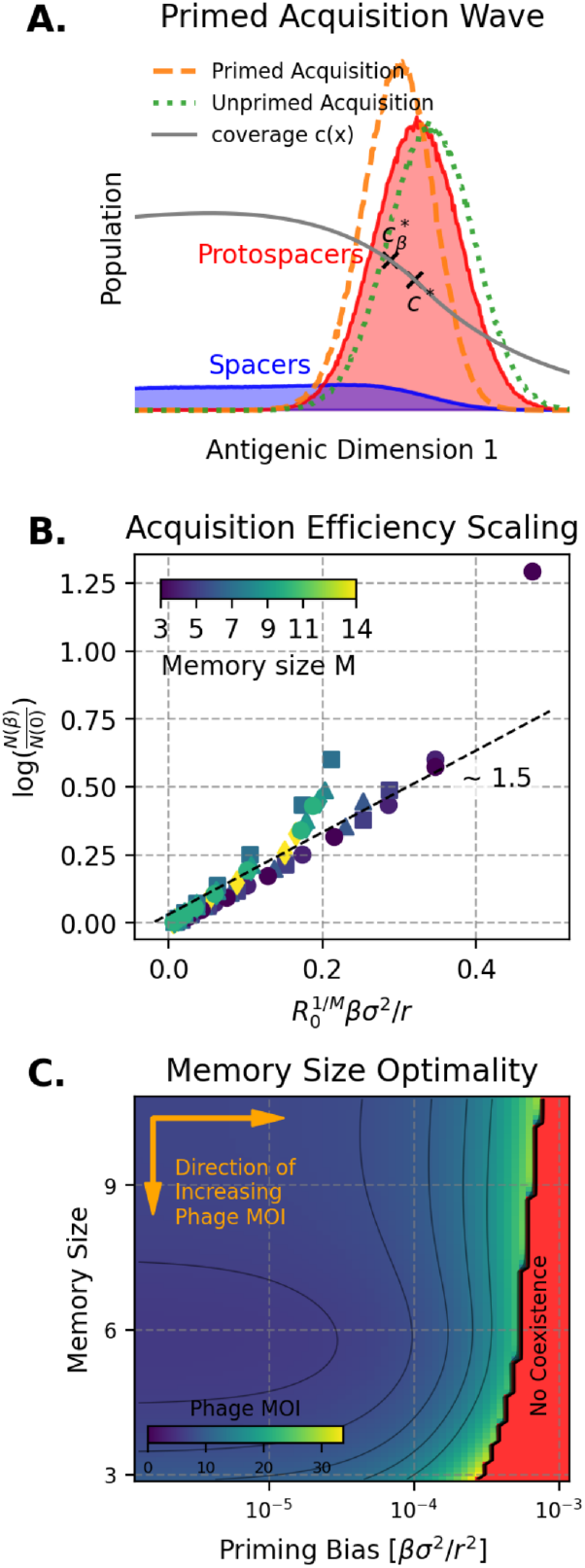
Primed acquisition. Simulations were done with *r* = 1, *R*_0_ = 100, *N*_*p*_ = 100, *α* = 0.01 and *D/r*^2^ ≈ 10^*−*5^.(A) Ac-quisition profile of primed and unprimed acquisition. Primed acquisition induces a delay in the phage protospacer acquisition probability. 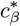 is the new average protospacer coverage after accounting for the delay. Unprimed acquisition is a theoretical complement of the primed case where bacteria try to predict future phage phenotypes. (see SI Fig. S6) (B) Scaling of relative phage MOI increase with priming. Assuming no change in acquisition rates, phage MOI increases exponentially with priming. Manually drawn line with 1.5 slope. (C) Coevolving Phage MOI heatmap to show memory size optimality given priming. Optimality effect are more pronounced at lower priming and goes away as priming bias increases. At high priming bias, the coevolving phage MOI is very high and the optimal number of memory increases. Note that the red region denotes regimes where the coevolving phage population is too large for a traveling wave solution.

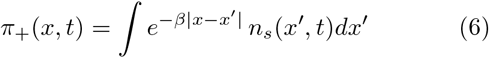

The acquisition profile for the whole spacer population depends on the multiplicity of each phenotype, depending on the current phage population: *P*_+_(*x, t*) ∝ *n*(*x, t*) *π*_+_(*x, t*) (Eq. 6). Under the assumption that our new protospacer population remains a normal distribution, we obtain a new acquisition profile with an antigenic delay of *βσ*^2^ in the traveling wave coordinate (Fig. 3A). It is important to note that this delay has an upper bound to the expected distance to the fittest individuals |*βσ*^2^| *< u*_*c*_ since coexistence requires an overlap between the acquisition probability *π*_+_ and the protospacer population distribution. Furthermore, within the previous assumption, it maintains all the previous constraints of the linear fitness regime. While it may seem counterintuitive that individual spacers exhibit a weaker acquisition bias compared to their cross-reactivity: 1*/β* ≪ *r*. It is important to recognize that the primed parameter is an averaged effect over all possible acquisitions, including naive. Although the presence of a primed spacer biases acquisition, bacteria can still engage in naïve spacer acquisition. In contrast, cross-reactivity is an intrinsic phenomenon of CRISPR immunity that cannot be turned off.

Finally, we obtain a new coverage profile in front of the spacer distribution. While the coverage at the center of the phage wave should remain at *c*^∗^ to maintain the zero growth condition. Priming induces a delay between the two waves (SI Sec. 4) and thus the coverage in front of the spacer wave must increase with priming. This leads to a different coverage at the leading edge of the spacer population:

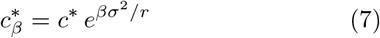

Then, using Eq. 6 and Eq. 7 and all the other constraints on *c*^∗^, we can get an estimate of the increase in phage MOI from priming (Eq. 8) (see SI Sec. 4A)

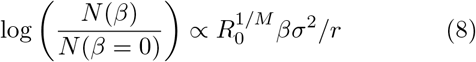

The most striking result of our theory and simulations in this section is the immediate increase in phage MOI once priming is introduced (see Fig. 3B). This indicates that across all memory sizes, the overall bacteria population performance worsens as the strength of the priming increases (assuming that acquisition rates *α* remain identical). Nevertheless, bacterial systems with larger memory sizes tend to perform better than smaller memory sizes as they retain more information since new information becomes rarer and cross-reactivity more important (see Fig. 3C). The advantage conferred by memory optimality also becomes less significant: the reduction in immune effectiveness caused by priming outweighs the benefits of memory retention.

Biologically, priming was thought to be beneficial because it increases the base acquisition rate of the bacterium for incoming primed spacers. While the underlying mechanisms have been described extensively, many sub-processes remain poorly understood. In our model, we show that a mere threefold increase in acquisition rate is enough to counteract the effect of primed acquisition over the range of priming values (see Fig. 3B, see also SI Fig. S5). In multiple lab experiments, priming bacteria seem to result in a 100-1000-fold increase in spacer acquisition [17], leading us to believe that the biological benefit of faster acquisition outweighs the evolutionary trade-offs introduced by priming. The net increase in maximum effective priming strength at larger memory sizes (Fig. 3C) might also become more relevant if evidence were found that specific spacers increased the priming bias independent of acquisition rates. The net effect of priming is also predicted to be more visible in CRISPR bacteria coevolving with slower-mutating phages since the range of visible difference due to priming increases: a slower-evolving protospacer population leads to a smaller increase in primed MOI.

### A. CRISPR Cassette Expansion

Most experiments on CRISPR-Cas adaptation show that spacer acquisition and loss often occur in bursts.

In particular, many studies report that the majority of spacers acquired during an experiment appear within only a few bacterial generations [2, 20, 28]. In contrast, spacer loss typically occurs on longer timescales, with evidence suggesting that deletions often remove spacers in blocks [51].

More generally, there are many more examples where the number of spacers in a CRISPR cassette must be transiently changed, yet the consequences of these fluctuations for coevolutionary dynamics remain poorly understood. For example, when CRISPR-Cas systems are encoded on mobile genetic elements [52], their presence can be dynamically regulated: these elements can be gained or lost by bacterial populations depending on environmental conditions [53]. Furthermore, strong experimental evidence indicates that horizontal gene transfer events often occur when the adaptive immune system is suppressed or temporarily inactivated [54].

While there are different methods for modulating and/or disabling CRISPR-Cas activity, such as Cas gene deletion, transcriptome insertion, etc. [55]. These observations illustrate a broader evolutionary principle: adaptive immunity confers long-term benefits, but there may be short-term benefits of modulating immune memory size.

To model the effects of a single expansion event, the total number of bacteria *N*_*b*_ will remain constant, with transient fluctuations that temporarily increase their cassette size (Fig. 4A). The idea is that bacteria will receive an excessive amount of new spacer for one generation while being disadvantaged for a short time (about 100 generations in our model to match typical bacteria recovery time) before returning to steady state. Immediately following an increased acquisition event, the individual memory size of each bacterium *M* (*t*) undergoes a recovery to its original or baseline memory capacity *M*_0_, which we model as an exponential recovery with recovery time *τ*_*m*_. The impact of each expansion event is governed by *γ/α*, which is the ratio between the increased acquisition compared to its baseline acquisition rate. Finally, for longer simulations with multiple events, we assume that those occur with a uniform time distribution with expected time between events to be longer than both the recovery timescale and escape timescale: *T*_HGT_ ≫ *τ*_*m*_, *T*_escape_. The escape timescale is the time required for the phage population to move beyond the range of cross-reactivity: *T*_escape_ = *v/r*.

**FIG. 4.**
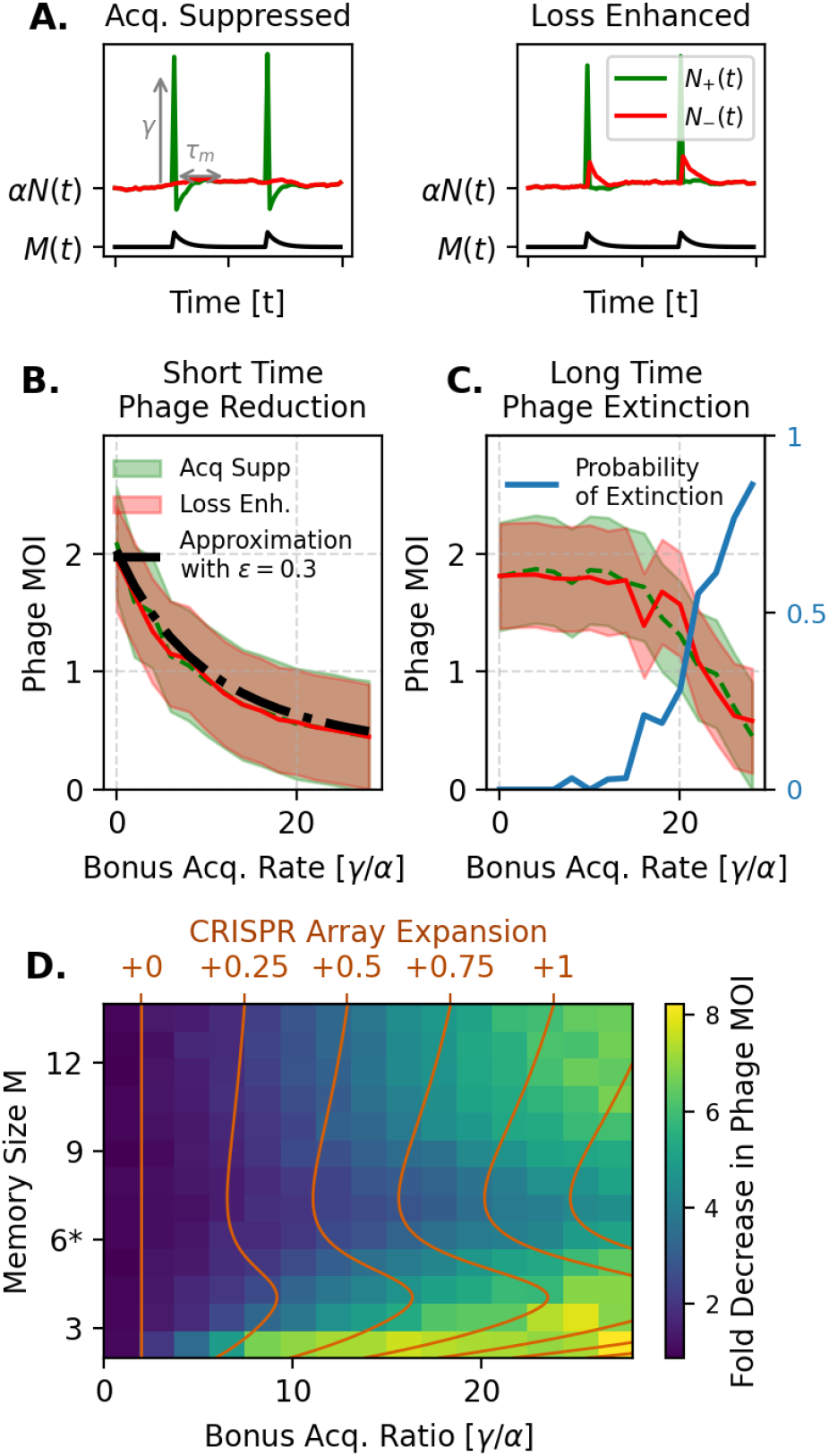
Memory Fluctuation plots where the memory size of bacteria fluctuates with time. Simulations were done with *r* = 1, *R*_0_ = 100, *N*_*p*_ = 100, *α* = 0.01, *D/r*^2^ ≈ 10^*−*6^ and *τ*_*m*_ = 100. (A) The two different schemes of memory recovery, where the tradeoff for short-term increase in memory size leads to a recovery time *τ*_*m*_ of either suppressed acquisition or increased spacer loss. (B) Simulations show that both scheme is equivalent in the short and long timescales. Short-term phage MOI decreases accordingly with bonus acquisition rate. Transient phage MOI were taken at *τ*_*c*_ = 20. (C) Longer simulations were taken at *T*_escape_ ∼ 150. As the fluctuation strength increases, the probability of extinction increases. (D) Fluctuation size depends on memory size; being at the optimal memory size (6*) does not lead to the most optimal decrease in phage MOI. The largest decrease in the transient MOI occurs when the memory size is small. Otherwise, the smallest increase occurs close to the optimal memory size

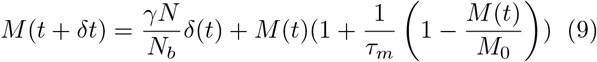

To get an estimate of the effect of immediately adding spacers to the system we can estimate the transient phage population by averaging out the fitness effects *N* (*t*) ≈ exp(∫ ⟨( *f* (*x, t*)⟩_*x*_*dt*)*N*_0_ where *N*_0_ is the original phage coexistence MOI and since we are in the linear fitness regime: ⟨*f* (*x, t*)⟩_*x*_ ≈ *∂*_*c*_*f* (*c*^∗^) |*c*_*δ*_(*t*) − *c*^∗^|, we only need to consider the effect of the increased memory perturbation at the start of the event. We used the previously derived methods from antigenic traveling wave theory to find the appropriate expression for the net perturbation of coverage in time:

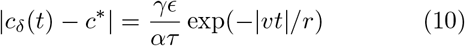

The *ϵ* term in Eq. 10 accounts for corrections arising from the spatial structure of the fluctuation. During the expansion event, most protospacers in the bulk of the population experience reduced infectivity and therefore go extinct, while only those located sufficiently far from the center of the traveling wave can persist. Finally, this leads us to a simple expression for the transient protospacer population (Eq. 11) with an implicit dependence on memory optimality coming from *∂*_*c*_*f* . Analytical results for longer timescales can also be derived (see SI Sec. 5B). Moreover, within the parameter regime considered here, the protospacer population rapidly returns to the coexistence state, and we will show that long-term dynamics are largely insensitive to the specific details of the recovery process.

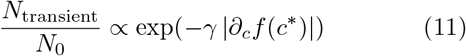

There are two extreme forms of recovery dynamics we can impose on the system. Following an increased acquisition event, the number of spacers acquired must be either suppressed or the amount of spacers lost must be enhanced for the system to return to its initial state. This is modeled by either a suppressed amount of acquired spacers *N*_+_(*t*) (Eq. 13) or a time-dependent enhanced number of spacers lost *N*_−_(*t*) (Eq. 12 and Fig. 4A). Those two forms of recovery dynamics correspond to different biological mechanisms associated with immune suppression, more specifically, whether the acquisition mechanism or the interference mechanism is considered. In either case, the complementary rate is kept at *αN* (*t*) so that only one component of the spacer turnover dynamics is perturbed in a given scenario.

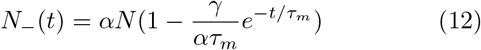

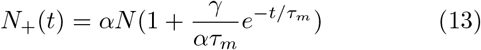

Overall, our simulation results indicate that cassette expansion primarily affects the short transient timescale of the evolutionary dynamics. In particular, its impact is limited to the time-to-effect *τ*_*c*_ = *u*_*c*_*/v*, which represents the delay before fitness changes propagate to the phage population at the leading edge of the distribution. When averaged over a longer period of time, such as the escape time *T*_escape_, the average phage population remains effectively unchanged regardless of the recovery scheme (see Fig. 4C). One reason for this robustness is that the magnitude of the fluctuations remains small even when individual expansion events temporarily increase spacer acquisition. Experimental observations show some support in this assumption: in laboratory studies [20, 28], although some individual cells acquired multiple spacers, most cells gained at most one spacer during the course of the experiment. This behavior is consistent with the average expansion rate observed in our simulations (see Fig. 4D). Furthermore, the strong selective filtering of spacer–protospacer interactions produces the same qualitative effect on the protospacer population as observed in our simulations (see Fig. 4C). As a result, when the expansion size is small, the resulting coexistence multiplicity of infection (MOI) remains very close to the unperturbed steady state, even if spacer deletions were to be permanent. This contrasts with the high expansion regime seen in experiments, where the expansion drives protospacer populations to extinction before the system transitions to a different evolutionary steady state.

However, memory optimality significantly alters the impact of cassette expansion during transient phases. In contrast to primed acquisition, smaller memory sizes are more effective at reducing the phage population than larger memory sizes, although both extremes outperform memory sizes near the optimal value (see Fig. 4D). While smaller cassettes are typically associated with higher baseline phage MOI, the addition of even a few spacers during expansion events produces a substantial reduction in protospacer abundance. Larger memory sizes also benefit from transient increases in spacer acquisition, but the resulting decrease in phage population is comparatively smaller. Populations with large memory sizes already operate far from the optimal trade-off point, and their larger coexisting phage populations might lead to the acquisition of many additional protospacers without a proportionally strong fitness gain. Analytically, this effect can be understood through the term |*∂*_*c*_*f*|, which represents the protospacer fitness sensitivity to coverage. This quantity is largest when the memory size *M* is kept small, thus leading to a larger decrease according to Eq. 11. Consequently, the largest cost of memory optimality arises when memory sizes are close to or slightly above the optimal value.

## IV. DISCUSSION

We have shown that while optimal memory size in CRISPR-Cas systems is maintained in a dynamical equilibrium environment, different evolutionary tradeoffs emerge when phage and bacteria are considered together as dynamic variables. In a statistical setting, the limited number of Cas interference proteins and their binding sensitivity suggest the existence of an optimal cassette size. Under dynamical coevolution, this optimality shifts depending on the presence of primed acquisition and cassette expansion.

Primed acquisition tends to favor larger cassette sizes. Although individual spacers acquired through priming may confer weaker protection, this lowers the expected coverage of the spacer memories, which leads to an increased benefit from maintaining more memory, thereby pushing the optimal memory size higher. Furthermore, the substantially higher acquisition rate observed in biological systems compensates for the reduced specificity.

In contrast, cassette expansion strongly benefits bacteria at smaller cassette sizes. When the goal is to drive phage populations to extinction rather than maintaining a smaller coexistence population, smaller increases in memory can yield a large selective difference for the bacterium. This is due in part to the fact that a small increase in coverage at low memory size can decrease the probability of infection substantially. This effect diminishes at larger memory sizes, where the relative benefit of an additional spacer becomes marginal.

In natural environments, bacteria usually have multiple cassettes in their genomes with different subtypes and cassette sizes [3]. Longer cassettes have been argued to better cover long-term coverage of common phage strains [12] and smaller memory size to rapidly acquire immune resistance to virulent strains [56]. Here, we suggest that the differences in evolutionary dynamics might also help explain the difference in cassette size depending on the eco-evolutionary constraints of the system. For phage-bacteria populations with longer coexistence time where phage and bacteria evolve gradually with one another, priming the spacer acquisition in exchange for increasing the acquisition is a logical evolutionary solution. In environments with highly diverse populations, competitive exclusion can be achieved more easily at the lower end of the memory sizes. Furthermore, the longterm cost of memory fluctuations seems to be minimal. Small fluctuations tend to revert to the traveling wave equilibrium, while large fluctuations are more likely to result in phage extinction. Thus, the system either self-restores or achieves its desired outcome depending on the fluctuation scale, without undermining long-term host fitness.

We have confirmed that the dependence of traveling wave dynamics on memory size holds both in one and two-dimensional systems. An important aspect not addressed in this work is the effect of priming and cassette expansion on phage speciation. In higher-dimensional systems, the phage population can spontaneously split into multiple subpopulations, a process heavily dependent on cross-reactivity. We hypothesize that both priming and memory fluctuations could affect the rate and likelihood of such splitting. However, speciation was beyond the scope of this study as it is a separate measure not directly tied to memory size, which could be an avenue for future research.

Furthermore, while we assume that all the different tradeoffs associated with priming and expansion share unified dynamics, a more ecologically realistic analysis should include phages with varying infectivity. For instance, separate phage populations with different infectivity might lead to differing extinction probabilities with more meaningful ecological consequences. This would allow for a more nuanced analysis of how different memory strategies perform under diverse selective pressures, potentially revealing more meaningful aspects of the tradeoffs.

Additionally, another limitation of our traveling wave framework is that we neglected the effect of oscillations and other periodic dynamics in the underlying predatorprey model. Those oscillations play an important role in understanding the effect of population collapse and runoff, particularly in the case of large memory fluctuations. While we assume that under expansion, the dynamical regime of linear fitness remains valid during fluctuations, this may not hold in more extreme scenarios where alternative traveling wave solutions could become relevant. Indeed, more complex oscillatory behavior in CRISPR-virus coevolution has also been predicted to appear due to non-linear dependence of the host immunity on the virus load [42], which our current model does not capture. This limitation arises from the simplifying assumptions built into the total population dynamics of the model, which favors a stable dynamical regime in favor of a detailed spatial description of our populations (see Supplementary Information, Section 5C).

Finally, a key component of our framework assumes all spacers are shared collectively across the population, even though we showed that this model could be used to explain agent-based simulation results that track individual spacer cassettes [27]. Nevertheless, the validity of this shared-memory approximation remains an open question and should be more critically evaluated by using empirical data from natural bacterial populations. While we hypothesize that our model is most applicable to populations in hot-spring communities, there is a strong lack of quantitative evidence to properly define the limits of our assumptions. Additionally, our dynamics are primarily driven by phages whose population dynamics remain difficult to characterize and notoriously hard to sequence.

Recent advances in single-cell sequencing and large-scale metagenomic analysis have provided unexpected insights into the communal nature of bacterial defense systems [57]. We hope that future experimental work leveraging these approaches can provide further insights into the role of horizontal gene transfer in spreading CRISPR elements and other immune components. Furthermore, a more detailed description of the interplay between individual and collective immunity could further extend our current simple model of memory size fluctuations.

## CODE AVAILABILITY

Code for this paper is available at this link.

## ACKNOWLEDGMENTS

This research was funded by the Natural Sciences and Engineering Research Council of Canada (Discovery Grant and RGPIN-2021). We acknowledge helpful discussions and comments from all the members of the Goyal and Zilman groups.

