## Supplementary material for "Dynamic Regulation of the Immune Repertoire of Bacteria": SI_Appendix

### 1. Fitness Model and Coevolution Calculations

**A. Traveling Wave Approximation.** We use the same ballistic approach to solve our traveling wave model, demonstrating that adding an acquisition rate does not fundamentally change the dynamics of the model and keeps the same approximations as Ref. (1). By first solving for the equilibrium solution of the spacer population, we assume that $N, N_b, M$ are all kept constant:

$$\partial_t n_s(x,t) = \alpha N \left( n(x,t) - \frac{n_s(x.t)}{M\,N_b} \right) \tag{1}$$

Solving Eq. 1, we get $n_s(x,t) = \alpha \int^t e^{-(t-t')/\tau}\, n(x,t')\, dt'$ where $\tau = MN_b/\alpha N$. The ballistic approximation for $n(x,t) \simeq N\delta(x - vt)$ then comes in assuming that $v, \sigma \ll r$. in the final solution.

$$\begin{aligned} n_s(x,t) &= \alpha \int^t e^{-\frac{t-t'}{\tau}}\, N\delta(x - vt)\, dt' \\ &= \frac{MN_b}{|v|\tau} \int^t e^{-\frac{t-x/v}{\tau}}\, N\delta(t' - \frac{x}{|v|})\, dt' \\ &= \frac{MN_b}{|v|\tau} e^{-\frac{vt-x}{v\tau}}\, H(vt - x) \end{aligned} \tag{2}$$

The final solution for the coverage $c(x) = \frac{1}{MN_b}\int e^{-\frac{|x-x'|}{r}} n_s(x',t)\, dx'$ is identical to the reference given a change of scaling in $\tau$. Ahead of the space wave, where $x > vt$, in the traveling frame of reference $u = x - vt$:

$$c(u) = \frac{1}{1 + \frac{v\tau}{r}} e^{-|u|/r} \tag{3}$$

Following from Ref. (1), we also have analytical solutions for the general velocity and spread of a linear fitness wave. Starting from linearizing $f(x,t) = f(c^*) + su$ where $s = |\partial_u f(u)|$:

$$\begin{aligned} \sigma &\approx (D/s)^{1/3} (24 \ln(N(Ds^2)^{1/2})^{1/6} \\ v &\approx D^{2/3} s^{1/3} (24 \ln(N(Ds^2)^{1/2})^{1/3} \end{aligned} \tag{4}$$

We also have a formula for the distance of the fittest population ahead of the wave: $u_c = s\sigma^4/4D$ (formula originally from Ref. (2)):

$$u_c \approx \frac{1}{4}(D/s)^{1/3}(24\ln(N(Ds^2)^{1/2})^{2/3} \tag{5}$$

Finally we can use $s = \partial_u f(u) = \partial_c f(c^*)\, \partial_u c(u)$ to find $s$. $\partial_u c(u)$ can be found using Eq. 3 but $\partial_c f(c)$ requires Eq.4 and Eq.5 of the main text:

$$\partial_c f(c) = \frac{M}{R_0} \frac{(1 + Q + (M-1)\frac{c}{1-c} Q)}{1 - c + McQ} \tag{6}$$

We find $c^*$ by finding the zero of the fitness function, then plugging into Eq.6 should give us $s$, which can then be used to find $v$ and $\sigma$ using Eq. 4. The constraints come from the fact that $s$ is defined by $\partial_c f(c)$, which depends on $v, \sigma$ from Eq.3. Therefore, all equations presented here define a linear fitness regime that fixes the coevolutionary protospacer population size $N$.

**B. Memory Optimization Calculation.** In this section, we adapt the methods and functions from Ref. (3) for finding the optimal memory size given $N_p$ constraints. In the previous work, there was also a threshold $p_c$, which is defined by the smallest number of complexes needed to successfully achieve interference; however, in this work, we take $p_c$ to be fixed at 10 for convenience. Furthermore, while the previous paper assumed a finite viral repertoire, we instead assume a constant bacterial immune coverage $c$, which is usually given by our traveling wave dynamics discussed in Sec.1. Let's start by considering the probability of infection:

$$\begin{aligned} P_{\text{inf}}(c) &= (1-c)^M + Mc(1-c)^M \sum_{p=0}^{p_c} \text{Binom}(N_p, p, 1/M) \\ &= (1-c)^M + Mc(1-c)^M\, Q(N_p, p_c, M) \end{aligned} \tag{7}$$

For simplicity, here is the definition of $Q$ used in both the main text and the rest of this supplementary:

$$Q = \sum_{p=0}^{p_c} \binom{N_p}{p} (\frac{1}{M})^p\, (1 - \frac{1}{M})^{N_p - p} \tag{8}$$

| Symbol | Description | Value |
|---|---|---|
| $\delta x$ | Simulation Length Scale | $10^{-3}$ bp |
| $N_b$ | Bacteria population | $10^4 \sim 10^6$ |
| $\delta t$ | Phage & bacteria generation time | $1$ |
| $M$ | Number of spacer per bacteria | $2 \sim 15$ |
| $R_0$ | Reproduction ratio of phage | $100$ |
| $N_p$ | Number of Cas proteins per bacteria | $10^2 \sim 10^3$ |
| $p_c$ | Cas protein activation threshold | $10$ |
| $\mu$ | Mutation rate | $1 \sim 10$ |
| $D = \frac{\mu < \delta^2 x >}{2}$ | Diffusion constant of mutation | $10^{-6} \sim 10^{-5}\ \delta^2 x/\delta t$ |
| $r$ | Cross-Reactivity radius | $1000\delta x$ |
| $\alpha$ | Spacer acquisition rate | $10^{-2}$ |
| $\beta$ | Priming strength | $10^{-3} \sim 10^{-1}$ |
| $\gamma$ | Memory size fluctuation strength | $10^{-4} \sim 10^{-1}$ |

**Table S1. Simulations values**

| Symbol | Description | Value |
|---|---|---|
| $\tau$ | Spacer persistence time | $10^1 \sim 10^2\ \delta t$ |
| $v$ | Phage adaptation velocity | $10^1 \sim 10^2\ \delta x/\delta t$ |
| $N$ | Mean phage coexistence population | $10^5 \sim 10^7$ |
| $\sigma^2$ | Standard deviation of phage population | $10^1 \sim 10^2\ \delta x$ |

**Table S2. Values from coexistence**

It serves to encode both the effect of $N_p$, the number of Cas interference proteins and $p_c$, the number of binding complexes required to achieve an immune response. Notably, $Q$ is independent of $c$ and is constant in the traveling wave regime.

At small coverage $c \ll 1$: $(1-c)^M \sim e^{-Mc}$ and $Mc(1-c)^M \sim 1 - e^{-Mc}$. Furthermore, given that $N_p$ is large compared to $M$, the binomial function can be approximated by a normal distribution. The variance of the normal distribution is $N_p(1/M)(1-1/M)$ can be approximated by $N_p/M$ given $M > 1$.

$$\sum_{p=0}^{p_c} \binom{N_p}{p} (\frac{1}{M})^p\ (1-\frac{1}{M})^{N_p - p} \approx \sum_{p=0}^{p_c} \mathcal{N}\left(\frac{N_p}{M}, \frac{N_p}{M}\right) \tag{9}$$

Eq.18 then becomes $P_{\text{inf}} = e^{-Mc} + (1-e^{-Mc}) \sum_{d=0}^{d_c} \mathcal{N}(\lambda^{-1}, \lambda^{-1})$. where $\lambda = M/N_p$. Finally, to find the optimal $\lambda^*$, we consider $P_{\text{surv}} = 1 - P_{\text{inf}}$, which has the same optimal memory size but allows us to simplify $1 - e^{-Mc} = 1 - e^{-\lambda N_p/c} \sim \lambda N_p/c$ given that $Mc \ll 1$. We then use the following approximation $\sum \mathcal{N}(\lambda^{-1}, \lambda^{-1}) \sim \int \frac{\lambda^{1/2}}{\sqrt{2\pi}} \exp(-z^2)\, dz$ to solve for the optimality condition on $\lambda$:

$$1 = \sum_{p=0}^{p_c} \mathcal{N}(\lambda^{-1}, \lambda^{-1}) \left( \frac{3}{2} - \lambda \frac{p^{-2} + \lambda^{-2}}{2} \right) \tag{10}$$

**C. Equilibrium Condition.** In this section, we solve for the equilibrium coverage $c^*$ for the traveling wave regime. While in practice, we will use numerical methods such as Newton's root-finding method to solve for $c^*$ based on the fitness function $f(c) = 0$. Here we will describe an analytical approximation for $c^*$ given the nonlinear element $Q$, which encodes most of the biochemical constraints of the bacteria.

Starting from eq. 3, we obtain our first condition for the coverage at the center of the phage traveling population $c^* = c(0) = (1 + v\tau/r)^{-1}$ where $c^*$ was originally defined for solving $f(c^*) = 0$. Another way of finding this value is through the fitness function from Eq. 6:

$$f(c) = \ln\left( R_0 (1-c)^M (1 - M\frac{c}{1-c} Q) \right) \tag{11}$$

There is no closed-form solution for $c^*$ given the nonlinear perturbation of $Q$, but we can expand around $Q$ since the probability of failure while having a specific spacer is small below optimal memory size: $Q << 1$. We first define a function $g(x)$ to solve for $c^*$ when $g(x) = 1/R_0$

$$\begin{aligned} g(x) &= (1-x)^M + Mx(1-x)^{M-1} Q) \\ &= \underbrace{(1-x)^{M-1}}_{(*)}\ \underbrace{(1 + (MQ-1)x)}_{(**)} \end{aligned} \tag{12}$$

Up to second order, let $c^* = c_0^* + Qc_1^* + \mathcal{O}(Q^2)$ where $c_0^* = 1 - R_0^{-1/M}$ is the base case when $Q = 0$ or the coverage center when a single spacer guarantees CRISPR interference. Plugging into the two separate parts of Eq. 12 such that $g(c^*) = g(c_0^* + Qc_1^*) \approx R_0^{-1}$:

$$\begin{aligned}(1)\quad (1-c^*)^{M-1} &= (1 - c_0^* - Qc_1^*)^{M-1} \\ &= (1 - (1 - R_0^{-1/M}) - Qc_1^*)^{M-1} \\ &\approx \left(R_0^{-\frac{M-1}{M}} - (M-1)QR_0^{-\frac{M-2}{M}} c_1^*\right)\end{aligned} \qquad [13]$$

Substituting $c^*$ and dropping quadratic order terms in (∗∗):

$$\begin{aligned}(2)\quad (1 + (MQ-1)c^*) &= (1 + (MQ-1)(c_0^* + Qc_1^*)) \\ &\approx \left(MQ(1 - R_0^{-1/M}) + R_0^{-1/M} - Qc_1^*\right)\end{aligned} \qquad [14]$$

Finally, we solve for the linear approximation of $g(x)$:

$$\begin{aligned}g(c^*) &\approx \left(R_0^{-\frac{M-1}{M}} - (M-1)QR_0^{-\frac{M-2}{M}} c_1^*\right)\left(MQ(1 - R_0^{-1/M}) + R_0^{-1/M} - Qc_1^*\right) \\ &\approx R_0^{-1} - (M - \cancel{1})QR_0^{-\frac{M-1}{M}} c_1^* + MQR_0^{-\frac{M-1}{M}} - MQR_0^{-1} - \cancel{QR_0^{-\frac{M-1}{M}} c_1^*} \\ &= R_0^{-1} + Q\left[-MR_0^{-1} + MR_0^{-\frac{M-1}{M}} - MR_0^{-\frac{M-1}{M}} c_1^*\right] + \mathcal{O}(Q^2)\end{aligned} \qquad [15]$$

Since the root of the fitness is when $g(c^*) = R_0^{-1}$, the $Q$-order term must vanish:

$$\begin{aligned}0 &= -MR_0^{-1} + MR_0^{-\frac{M-1}{M}} - MR_0^{-\frac{M-1}{M}} c_1^* \\ c_1^* R_0^{1/M} &= R_0^{1/M} - 1 \\ c_1^* &= 1 - R_0^{-1/M} = c_0^*\end{aligned} \qquad [16]$$

The final result is:

$$c^* = c_0^*(1 + Q) \qquad [17]$$

This approximation is accurate when $Q$ is small, which is true when $M < M_{\text{optimal}}$ (see Fig. S1B). During numerical analysis and simulation initialization, we find $c^*$ using Newton's root-finding method since it gives us a more accurate solution for $c^*$ outside the validity range of the previous estimate.

Finally, we considered the case where a single effective spacer did not confer full immune coverage and where any higher number of immune spacers would guarantee protection. Alternatively, a full infection probability can be computed by taking into account the probability of all possible numbers of spacers up to $M$:

$$P_{\text{inf}}(c) = \sum_{i=0}^{M} \text{Binom}(i, M, c) \sum_{p=0}^{p_c} \text{Binom}(N_p, p, i/M) \qquad [18]$$

In Fig. S1A, we compare the infection probability calculated using the full set of effective spacers versus using only a single effective spacer for a given value of coverage. At small memory sizes up to $M_{\text{optimal}}$, the two approaches yield nearly identical total probabilities across all coverage values. However, when looking at longer memory sizes, the difference between the two formulations becomes more evident, with the maximum deviation occurring at around $c^* \approx 0.1$ under our usual simulation parameters ($N_p, dc = 100, 10$). Notably, for certain $c^*$ values, the local memory optimum in the single effective spacer formulation becomes a global minimum across all memory sizes for the full infection probability. Under our typical parameters, this discrepancy effectively caps the usable memory size at $M \leq 15$ in general simulations, and for primed acquisition, where the expected coverage is lower, we further restrict it to $M \leq 12$, placing us near the start of the infection plateau.

In summary, using a single effective spacer preserves analytical tractability while maintaining the same $M_{\text{optimal}}$ as the full formulation. To remain consistent with theory, we adopted the single-spacer formulation throughout our simulations, but limited the maximum memory size to remain within the regime where both formulations yield comparable results. In Fig. S1C, we illustrate the resulting differences in phage MOI for 2D simulations. This extension to higher dimensions is motivated by theoretical prediction suggesting that the 1D equations also extend to 2D along the principal axis of movement of wave propagation (1). Our results confirm that the 2D simulations are consistent with the expected multiplicity of infection (MOI) as predicted by our fitness model.

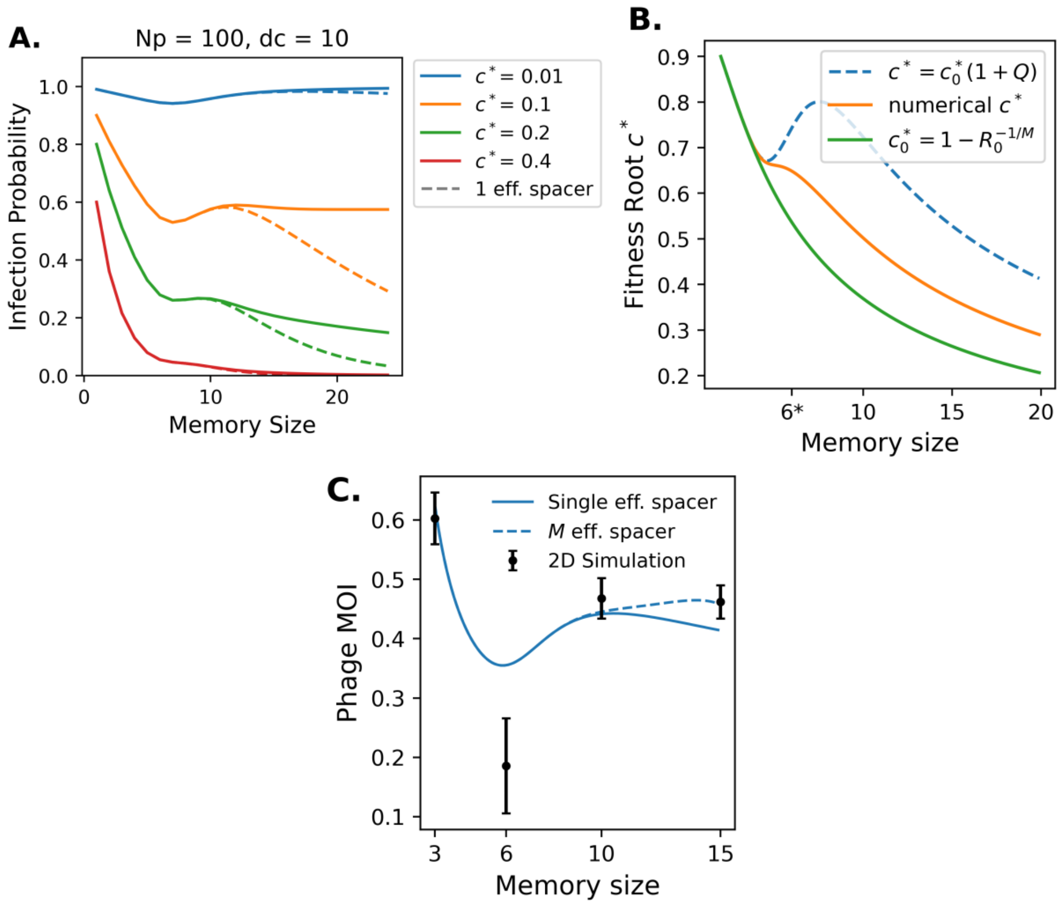


**Fig. S1.** Coexistence parameters of traveling waves. (A) Difference in probability of infection when only the effect of one effective spacer is considered vs. $M$ effective spacers. This assumption on the probability of infection is valid when the memory size is smaller than optimal: $M < M_{\text{optimal}}$ for all ranges of coverage. Outside this range, the assumption is still valid at very high ($> 0.4$) or very low coverage ($< 0.05$). (B) Parameter range for $c^*$, the coverage at the wave center, and the zero of the fitness function for $N_p, d_c = 100, 10$. While coverage usually decreases with memory size, we avoid the region of invalidity for the probability of infection by not simulating below $M = 15$. (C) 2D simulations of traveling wave, simulations were performed with $N_b = 10^5$, $\mu = 10$, $\alpha = 0.1$ and $N_p, p_c = 100, 10$. Optimal memory size is maintained for waves in 2D. The difference between one and multiple effective spacers is shown: the discrepancy is only significant when $M > 12$.

### 2. Material and Methods

The simulation procedure, in both 1D and 2D, consists of 3 distinct steps in order: (1) phage mutation, (2) coverage/selection, and (3) spacer Acquisition/Loss. In 1D simulations, the populations were implemented using `numpy.array` while in 2D, we used `scipy.sparse` to reduce memory usage and computational complexity. Finally, to accelerate computation in 2D, there is usually multi-threading involved. Those parallelizations were done using `joblib`. In particular, during the coverage calculation step, we relied on optimized matrix operations using `numpy.dot()`.

**A. Phage Mutation.** For each nonzero location of phage density location $n(x)$, we first estimate the total number of phage mutations per antigenic location by sampling from a binomial distribution with $n(x,t)$ tails and probability distribution $p = (1 - e^{-\mu\,\delta t})$. The expected number of protospacers undergoing mutations is $n(x,t) * p$. For each protospacer undergoing mutation, the number of mutation jumps is drawn from a Poisson distribution with mean $\mu\,\delta t$. Finally, each mutation step corresponds to a jump of random length drawn from a Gamma distribution with shape parameter 20 and mean $2\,\delta x$. The final position of each protospacer will be determined by the sum of all the jumps. For 2D simulations, each jump involves sampling an additional angle and direction from a uniform distribution in $[0, 2\pi)$. For multi-threaded simulations, after the number of mutants at each nonzero location is determined, each thread independently computes the mutations and resulting position of a subset of initial positions for which we have determined that a mutation must have occurred. The results are then aggregated into a new sparse matrix representing the updated phage density.

**B. Phage Fitness and Bacteria coverage.** For 1D simulations, the coverage $c(x,t)$ and the acquisition probability kernel $p_+(x,t)$ are both just computed using `scipy.signal.convolve`. The fitness $f(x,t)$ is evaluated according to Eq.6 in the main text. Phage selection is applied by sampling from a Poisson distribution with mean $(1 + f(x,t)\,\delta t) \cdot n(x,t)$. Sometimes, fitness values must be normalized to have zero mean by centering the fitness $f(x,t) - < f(x,t) >$ where $< f(x,t) >= \sum_x f(x,t)\,n(x,t)/N(t)$ is the average fitness of the phage population. This normalization was applied, for example, in Fig. 3A of the main text.

In 2D simulations, the coverage is directly performed through element-wise operations on sparse matrices. First, a dictionary of precomputed exponential values was initialized for all pairwise index separation distances $d(i,j) = \sqrt{i^2 + j^2}$ within a 4000x4000 convolution window. Each dictionary entry uses separation distance $d(i,j)$ as the key and stores the corresponding value $\exp(-d(i,j)/r)$. Recomputing these values avoids repeated exponential calculations and significantly accelerates convolution. Furthermore, since both matrices $n_s$ and $n$ are sparse, we have found that element-wise coverage still seems to be the fastest method over using FFTs to convolve subsets of the domain.

The coverage is then performed in parallel using a dot product. Each nonzero location of $n_s(i,j)$ (sparse matrix form of $n_s(x,t)$) is batched and distributed to each thread. Each thread computes a dot product using `numpy.dot()`, summing the contributions of phage density at positions within the convolution window. Specifically, for each pair of indices for each batch and from nonzero positions in $n_s(i,j)$, the index separation distance is calculated. If the distance falls within the precomputed convolution window, the corresponding dictionary value is added to the coverage sum for that location. While 2D primed acquisition was not shown in this paper, the single spacer acquisition kernel, $\pi_+$, is calculated similarly by simply replacing the precomputed dictionary to be $\exp(-\beta\,d(i,j))$. The full acquisition kernel $p_+$, is computed after the selection step by performing a multiplication with the resulting $n(x,t)$ post-selection. This also ensures ordering consistency as $n_s(x,t)$ is unchanged during coverage computation and phage selection.

**C. Spacer Acquisition and Loss.** Spacer acquisition will be done by first adding to the spacer population the expected number of new protospacers $N_+\,p_+(x,t)$ rounded to the lowest integer at each location. To make sure the total number of spacers acquired is exactly $N_+$, any remaining spacers will be assigned by sampling from the phage population with probability $p_+(x,t)$. Because the phage population is typically the largest quantity within our simulations, we believe this is a reasonable use of the mean approximation. This procedure remains valid in a multi-threaded context as well.

Spacer loss, on the other hand, is implemented by randomly sampling $N_-$ spacers from the current phage population $n_s(x,t)$ with each spacer having an equal probability of being lost. In the multi-threaded context, we partition $n_s(x,t)$ again into separate batches, and thread independently sample a portion of the $N_-$ losses in parallel. Any inconsistencies, such as negative spacer counts, are corrected after sampling. Although this parallelized loss method introduces slight deviations in the variance of the loss distribution compared to a serial implementation, we consider the faster simulation time to be a worthwhile tradeoff.

**D. Simulation Initialization and Analysis.** To initialize the simulation, we first compute the traveling wave parameters $N, \sigma, v, u_c, \tau$ from the traveling wave conditions (Eq.7-8 of the main text). The phage population $n(x,t)$ is initialized as a normal distribution with standard deviation $\sigma$, centered at $x = \beta\sigma^2$ with total size $N$. The spacer population $n_s(x,t)$ is initialized by numerically computing $\sum_t \alpha n(x,t')e^{-(t-t')/\tau}$ and then placing $N_b$ bacteria accordingly, such that the head of the distribution is positioned at $x = 0$. If no stable traveling wave solution was found, we proceed with the closest approximation based on Eq.7 (Main Text). For 2D simulations, both populations are extended along the transverse axis $y$ using a marginalized probability distribution with standard deviation $1.66\sigma$ and mean at $y = 0$. To track our 2D traveling distributions, we use `sklearn.mixture.BayesianGaussianMixture` to fit a probability distribution for our phage population. Phage runoff is defined as a situation where the current phage population exceeds $10\times$ the expected steady state phage population (outside of fluctuation simulations), and phage extinction is defined when the phage population drops below 1.

### 3. Embedding Details and Experimental Observations

**A. Agent-Based Simulation Embedding.** To construct the embedding, we first defined a shared sequence space containing all unique spacer and protospacer sequences present in the dataset. When applying this procedure to experimental data, protospacer-adjacent motif (PAM) sequences were excluded prior to analysis. We then computed a pairwise distance matrix over all found sequences. For binary vector simulation data, distances were calculated using the Hamming distance, while for experimental sequence data, we used the minimum number of nucleotide substitutions between sequences.

The resulting distance matrix was subsequently used as input to a multidimensional scaling (MDS) procedure in order to obtain a low-dimensional representation of the sequence space. Specifically, we applied the metric MDS implementation available in the `scikit-learn` library with the `metric=True` option to generate the final embedded map.

As shown in Fig. S2A, the embedded data clearly separates into two distinct lineages. To identify the corresponding spacer clusters, we applied $k$-means clustering to the embedded coordinates of all unique spacer sequences. To track the temporal trajectory of individual lineages, we then employed a random walk model similar to those previously used to describe the antigenic evolution of influenza viruses.

$$x_t = x_{t-1} + \mu \Delta t + \sqrt{\Delta t}\eta \text{ where } \eta \sim N(0, \Sigma) \quad [19]$$

In Eq. 19, $\mu$ and $\Sigma$ denote the drift vector and covariance matrix of the noise term, respectively. These parameters were estimated from the observed temporal evolution of the lineage centroids $\hat{x}_t$ in the data. Specifically, we computed

$$\hat{\mu} = \langle \Delta_t \rangle, \qquad \hat{\Sigma} = \mathrm{Cov}(\Delta_t),$$

where

$$\Delta_t = \frac{\hat{x}_t - \hat{x}_{t-1}}{\Delta t}.$$

In addition, we account for observational noise at each time point by estimating the dispersion of individual observations $y_t$ around the corresponding centroid $\hat{x}_t$, measured independently along each embedding axis.

$$y_t = \hat{x}_t + \theta \text{ where } \theta \sim N(0, R) \quad [20]$$

Finally, we applied a Kalman smoothing filter (4) to the inferred trajectories in order to reduce observational noise. The Kalman smoother estimates the most likely trajectory by combining information from neighboring time points within a temporal window, weighting observations according to the inferred noise covariance. To qualitatively assess the dimensionality required for the embedding, we partitioned the simulation data into two subsets. Specifically, 90% of the time points were used to fit the model, while the remaining 10% of the final time points were reserved for out-of-sample prediction. We first evaluated the mean root-mean-square (RMS) fitting error computed over the training portion of the dataset (the first 90% of time points):

$$\mathrm{rms}(d) = \sqrt{\frac{1}{N_t} \sum (y_t(d) - x_t(d))^2} \quad [21]$$

We first observe that when the two lineages are not separated, the RMS fitting error is large for a one-dimensional embedding but rapidly decreases and settles into the expected embedding dimensionality error. In contrast, when the dataset is partitioned into two distinct lineages, the fit becomes most accurate in a one-dimensional embedding, reducing the RMS distance by approximately an order of magnitude compared to all lineages combined (see Fiq. S2D).

We further evaluated predictive performance using the Mahalanobis distance (5) between the predicted trajectory and the held-out dataset (see Fig. S2E). This metric quantifies the number of standard deviations by which the prediction deviates from the model, accounting for the covariance structure of the inferred dynamics. For embeddings with dimensionality greater than two, the observed deviations exceed the expected $\sqrt{d}$ scaling across all lineages between 1 and 3 dimensions. Together, these results suggest that a low-dimensional embedding ($d \lesssim 2$) provides the most appropriate representation of the lineage dynamics.

Finally, the poorer predictive performance observed in higher-dimensional embeddings can be understood by examining the inferred drift and covariance parameters (see Fig. S2C). A larger drift term generally improves predictive power, as it implies a more directed trajectory and therefore greater certainty about the future position of the lineage. In contrast, a larger covariance indicates that the dynamics are dominated by stochastic fluctuations, reducing the embedding's ability to predict future states.

Examining the inferred parameters as a function of embedding dimension reveals that, when lineages are not separated, the estimated drift parameter in one dimension exhibits a large inference error. Partitioning the dataset into distinct lineages substantially reduces this error (see Fig. S2F-H). Moreover, the inferred covariance increases systematically with embedding dimension. This increase indicates that higher-dimensional embeddings attribute a larger fraction of the dynamics to stochastic motion, which in turn explains the observed decrease in predictive accuracy.

**B. Experiments.** While we do not yet have direct experimental evidence of a coevolutionary traveling-wave regime, our analysis of the spacer dataset from Paez-Espino et al. (2013) (6) suggests that several spacer labels may correspond to the early stages of such dynamics. One limitation of this dataset is its relatively short duration: the experiment spans only $\sim 100$ bacterial generations, as the coevolution experiment was conducted over a period of 15 days. This limited time length may be insufficient to fully resolve the emergence of traveling-wave trajectories.

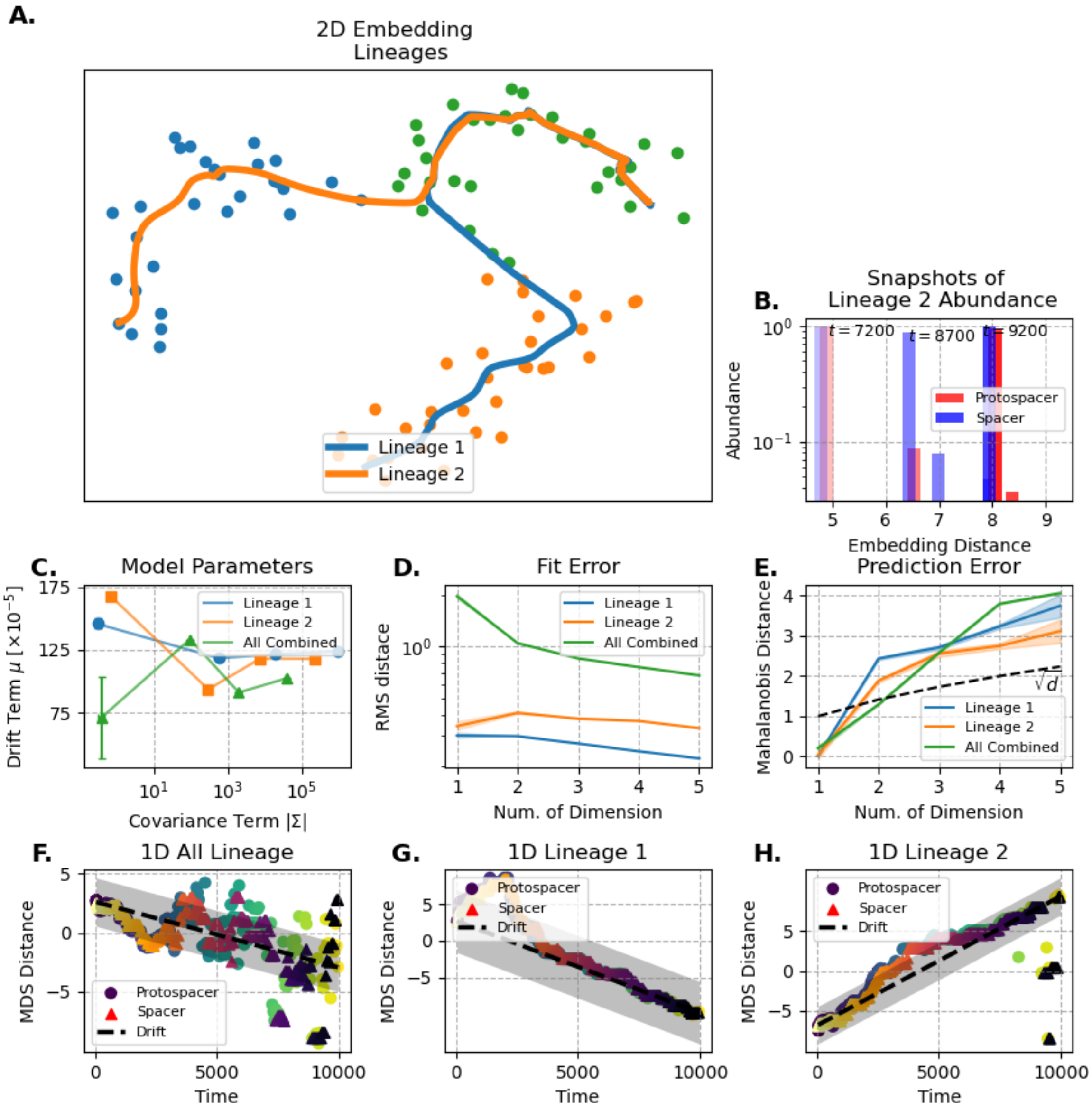


**Fig. S2.** Details of Agent-based Simulation Embedding. (A) Lineages found by fitting a random walk trajectory. Clusters were found using kmeans. (B) Snapshots of phage/protospacer population at any given time over a given lineage. Note how most population are only within one substitution of each other. (C) Drift vs Covariance (determinant of covariance matrix) parameters of the random walk model. Each line follows an increasing dimension from 1 to 4 from left to right. Note how the drift parameter is inaccurate when all lineages are combined. (D) Fit/Model error over 90% of timepoints used to fit the random walk. Note how the jump from 1D to 2D significantly changes the error trend of embedded lineages. (E) Prediction error onto last 10% of timepoints used for validation. Note that lower dimensional embeddding are significantly more predictive even when accounting for the $\sqrt{d}$ scaling. (F-H) 1D Embeddings. While the 1D embedding of all lineages combined fails, 1D embedding works when lineages are separated.

Longer coevolution experiments exist, such as Paez-Espino et al. 2015 (270 days) (7) and Burstein et al. 2016 (3 years) (8). However, these studies lack the temporal resolution required to track evolutionary trajectories in detail, as they contain only 14 and 6 sampled time points, respectively. Consequently, substantial evolutionary change occurs between successive observations, making it difficult to reconstruct continuous traveling-wave dynamics.

We therefore hypothesize that an experiment designed to balance both duration and temporal resolution could reveal such trajectories. One possible approach would be to perform a longer coevolution experiment using *S. thermophilus* in milk culture, sequenced at a depth comparable to that of Paez-Espino et al. 2013, but extended for approximately 5–10 times longer. Using previously inferred parameters $R_0 \sim 100$, $D/r^2 \sim 10^{-6}$, and $M = 1.5 \pm 0.5$ (estimated from data) from Ref. (1), we can estimate the expected number of generations between successive lineages split:

$$t_{persist} = \frac{r^2}{4D} R_0^{-2/M} \approx 500 \text{ gen.} \qquad [22]$$

Finally, our labeling procedure differs slightly from the analysis performed in Bonsma-Fisher et al. (9). To ensure that no evolutionary information was lost due to overly aggressive clustering, we re-labeled the previously extracted spacers using a combination of $k$-mer distance (10) and DBSCAN clustering. The $k$-mer distance metric was first used to rapidly partition the dataset into ranked clusters. We then examined the distribution of pairwise $k$-mer distances to determine an appropriate cutoff threshold for the DBSCAN algorithm. This threshold was chosen to avoid producing clusters that were too small, which could otherwise exclude spacers that had drifted slightly from their original cluster over time.Based on our inferred parameters $(D/r^2 \sim 10^{-6})$ , the expected drift length over $\sim 100$ generations is approximately 1-2 nucleotide distance. The clustering threshold was therefore selected such that spacers within this evolutionary drift range would remain grouped within the same cluster (see Fig. S3B).

Finally, our modeling indicates that both $k$-mer and Hamming distance metrics preserve the same underlying clusters. This consistency can be observed in the aggregation of labels within the multidimensional scaling (MDS) embedding (see Fig. S3A): although the labels were assigned using $k$-mer distances, the embedding itself was constructed using Hamming distances, yet the same clusters remain separated.

Evidence that the Paez-Espino et al. 2013 coevolutionary dataset may contain signatures of an antigenic regime can be obtained by examining the phylogeny of individual spacer clusters (see Fig. S5). Some clusters (e.g., labels 0 and 1) exhibit phylogenetic structures consistent with low cross-reactivity, while others (e.g., labels 6 and 12) display a chronological ordering along their branches that may be compatible with traveling-wave-like dynamics. However, because the experiment spans a relatively short evolutionary timescale, we cannot determine whether the embedding fully captures an established regime or whether the traveling-wave dynamics have not yet emerged for most clusters (see Fig. S4). Finally, clusters 18 and 19 appear only within a single sampled time point and therefore likely correspond to spacer lineages that have not yet diversified.

Finally, our analysis reveals that *Strep. thermophilus* exhibits a highly peaked spacer distribution in between each sampled time point, consistent with the predictions of the antigenic wave model when integrated between time points. In particular, the most abundant spacer is typically several orders of magnitude more frequent than its nearest mutant variant. A visual best-fit comparison suggests an effective antigenic mutation rate of approximately $D/r^2 \sim 10^{-6}$, which is consistent with experimentally reported rates of CRISPR escape mutation accumulation in the phage genome from the original study (6). We did not attempt to compare adaptation velocities, as the limited temporal resolution of the dataset and the substantial variance between sampled time points make such estimates unreliable.

### 4. Primed and unprimed acquisition

Primed acquisition is a shift of the acquisition probability kernel defined in Eq.4 of the main text. Here, we will derive the main results starting from the acquisition probability kernel We start with Eq. 2 to derived a modified version of Eq. 3 for our single spacer acquisition kernel $\pi_+(x,t) \propto \int e^{-\beta|x-x'|} n_s(x',t)\, dx'$. Since we will normalize this probability distribution afterwards, we only care about the shape of our acquisition kernel ahead of the traveling wave.

$$\pi_+(u) \propto e^{-\beta|u|} \qquad [23]$$

Finally to obtain the final normalized acquisition probability $p_+(u) = \pi(u)n(u)/\int \pi(u)n(u)\, du$. We solve for the quadratic term and drop all higher orders in $u$.

$$\begin{aligned} \ln p_+(x,t) &\approx -\beta u - \frac{u^2}{2\sigma^2} + \mathcal{O}(1) \\ &\approx -\beta u - \frac{u^2}{2\sigma^2} + \beta\sigma^2 - \beta\sigma^2 + \mathcal{O}(1) \\ &\approx -\frac{1}{2\sigma^2}(u^2 + 2u\beta\sigma^2 + \beta^2\sigma^4) + \mathcal{O}(1) \\ &\approx \frac{-(u+\beta\sigma^2)^2}{2\sigma^2} + \mathcal{O}(1) \end{aligned} \qquad [24]$$

In Eq. 24 on the second line, in order to complete the square, we assumed that $\beta\sigma^2$ remains constant and hence the variance in $p_+(x,t)$ and $n(x,t)$ remains similar. This is true largely because both wave velocity and variance remain largely dependent

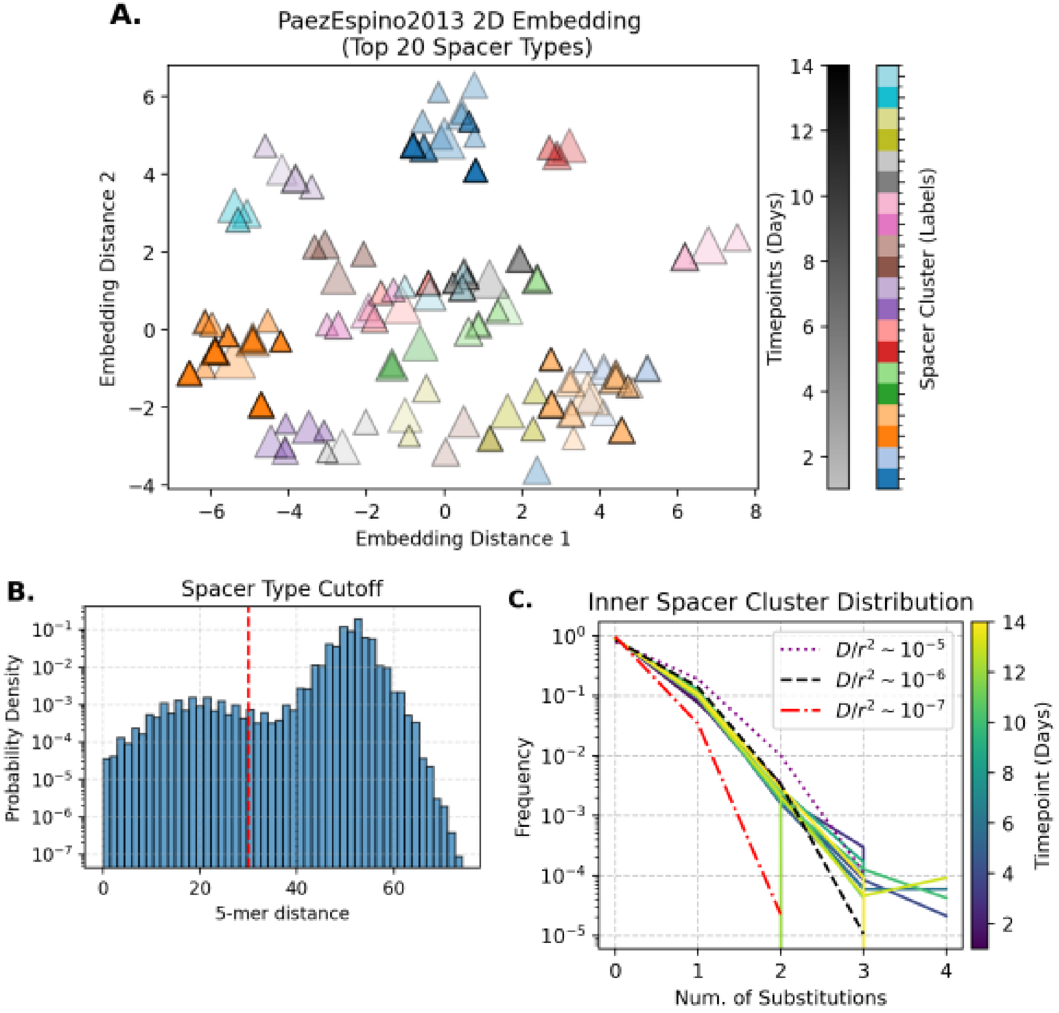


**Fig. S3.** Embedding details for embedding and travelling wave analysis of Paez-Espino 2013 (6). (A) 2D embedding of top 20 most abundant spacer labels. All spacer sequences were found and labelled into clusters using k-mer metric then the embedding was found by computing num. of minimun substitution over all spacers in the top 20 labels. Notice how most of the clusters are preserved from one metric to another. (B) K-mer distance cutoff used to compute cluster. The cutoff needs to be larger than the 1-2 substitution expected between timepoints. (C) Estimates of the number of substitution within cluster found between timepoints compared to traveling wave theory. visual estimates put the mutation rate $D/r^2$ at approximately $10^{-6}$.

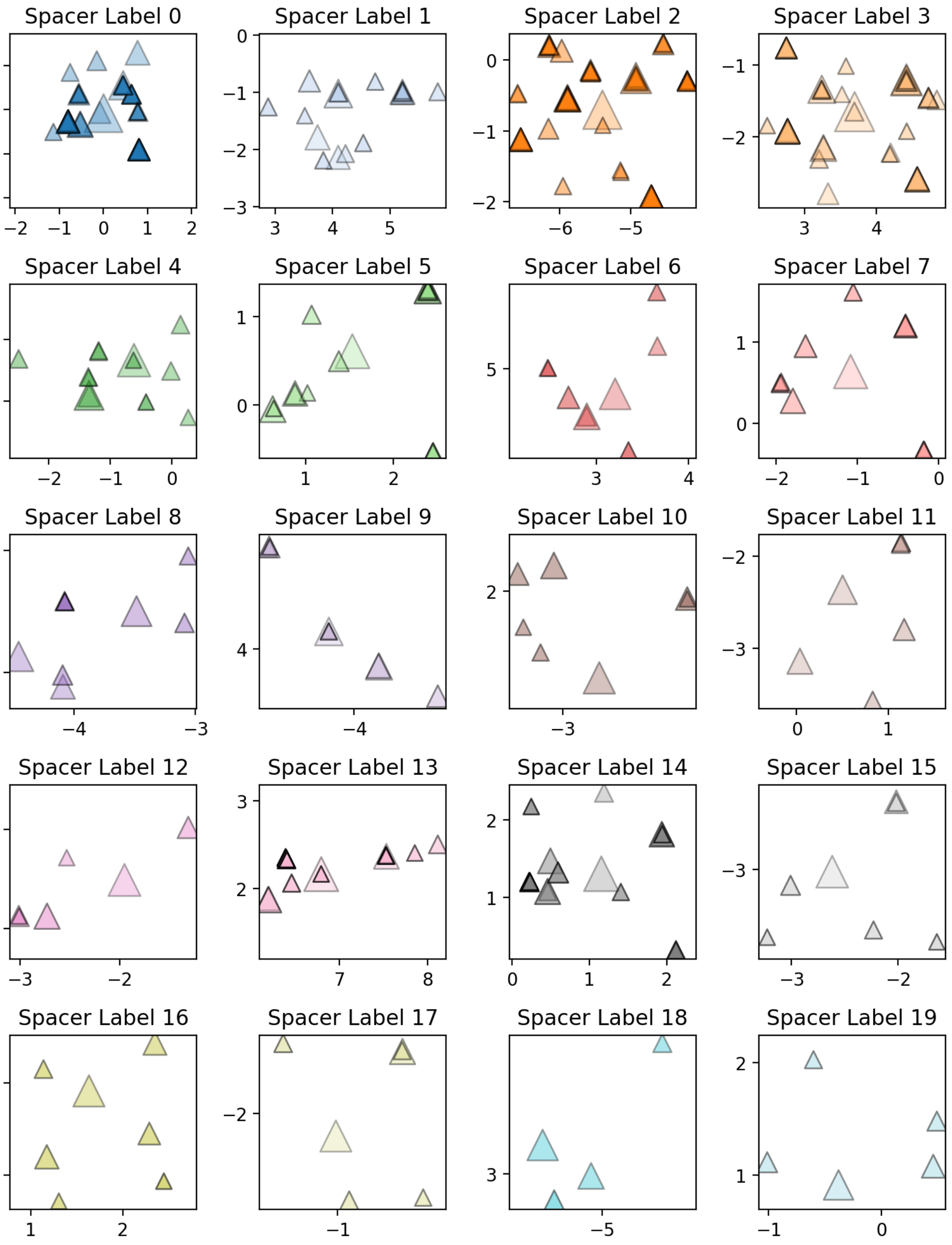


**Fig. S4.** 2D embedding of the 20 most abundant spacer clusters. Marker opacity indicates time (earlier time points are more transparent), and marker size scales with log abundance.

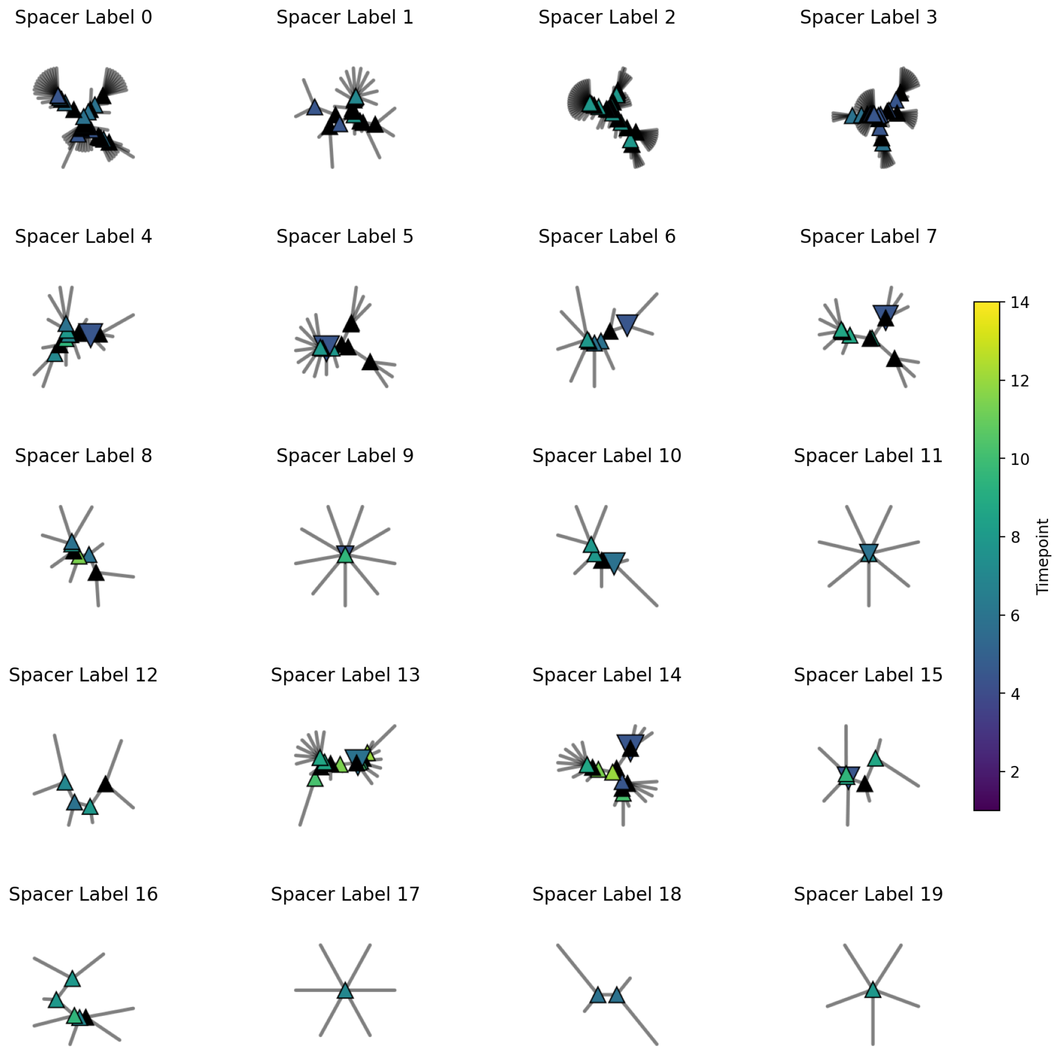


**Fig. S5.** Unrooted phylogenetic tree of spacers. Labels 6 and 12 are consistent with the embedding criteria described in the main text, whereas the remaining labels lack sufficient temporal resolution to support a reliable embedding

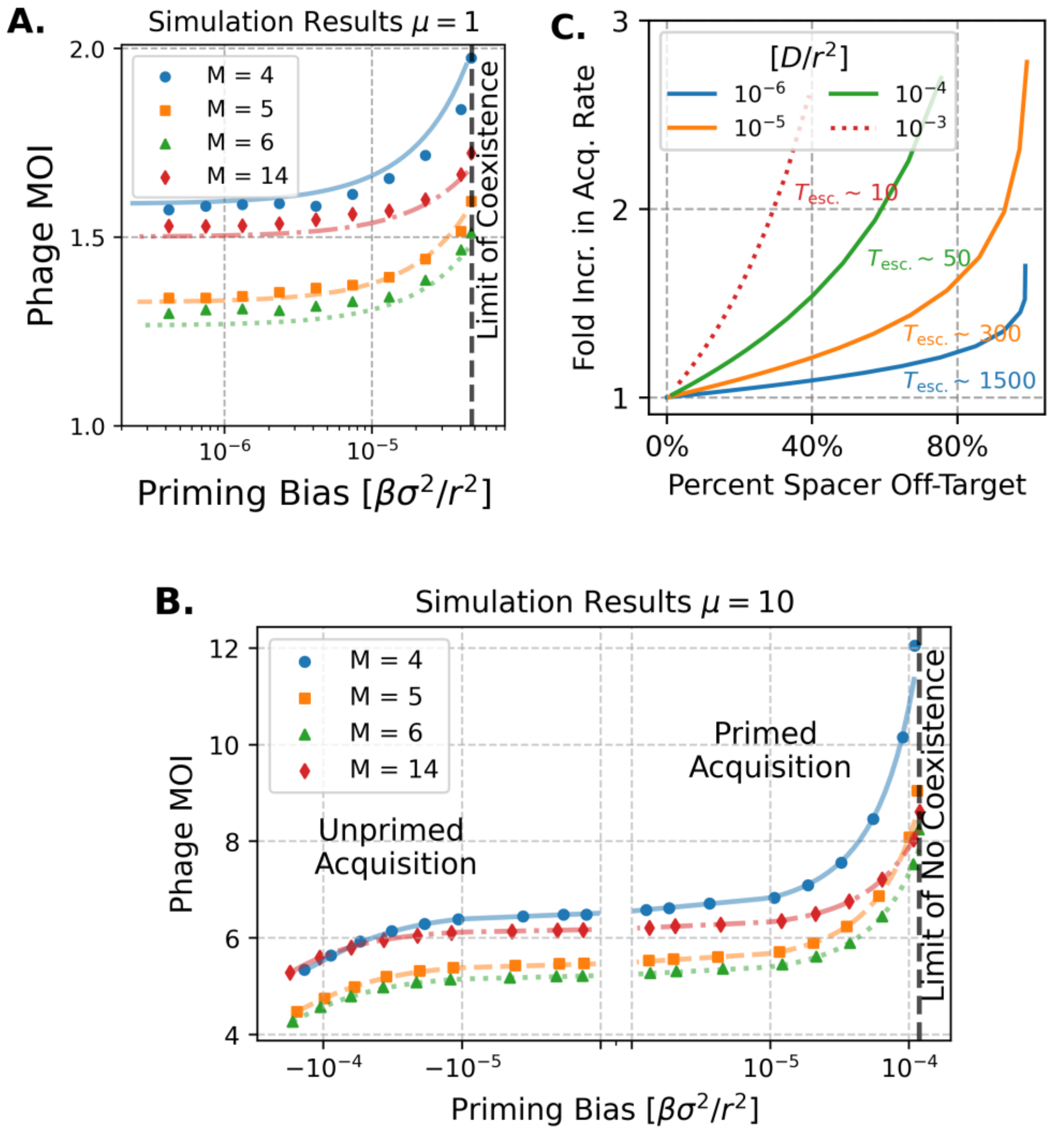


**Fig. S6.** Phage MOI scaling with priming: theory and simulation. (A) Theoretical priming scaling at low mutation rate, corresponding to $D/r^2 \approx 10^{-6}$. The phage MOI changes only weakly. The coexistence boundary occurs where the traveling-wave solution no longer satisfies the model constraints. (B) Simulation results for $D/r^2 = 10^{-5}$. Unprimed acquisition corresponds to the negative-$\beta$ extension of the primed-acquisition theory and has little effect on phage MOI. (C) Effective increase in acquisition rate as a function of mutation rate. At higher mutation rates, a larger acquisition rate is required to compensate for the same priming strength. The x-axis (priming strength) is rescaled to represent the fraction of spacers acquired outside the naïve acquisition distribution.

on mutation rate $\mu$ and weakly dependent on the number of phages $N$ from Eq. 7 of the main text. Finally, the $-\beta\sigma^2$ terms get absorbed in the $\mathcal{O}(1)$ term since this probability is normalized.

We thus obtain an approximation of the acquisition profile $p_+(u) \sim n(u+\beta\sigma^2)/N$, which is just a shift in the traveling frame depending on the priming strength $\beta$. This approximation is limited by the support of the phage population $n(x)$, which is around a distance $u_c = s\sigma^4/4D$ (see Eq. 5) around the center $u = 0$, thus giving us a limit of $|\beta\sigma^2| \leq u_c$. Combining this delay with eq. 3 gives us the expression for the shift in coverage due to priming:

$$c^*_\beta = c^*\, e^{\frac{\beta\sigma^2}{r}} \tag{25}$$

**A. MOI Scaling Behavior.** To obtain the scaling, we need to first compute $v\tau$ compared to $c^*$ from eq.3 (at the tip of the wave where $u = 0$):

$$v\tau = r(\frac{1}{c^*} - 1) \tag{26}$$

To obtain the required increase in acquisition rates, we compare $v\tau$ using eq. 25 to obtain our final expression. Finally, to find the equivalent increase in acquisition rates, we can compare $v\tau$. Since $v$ is again largely dependent on $\mu$ and weakly dependent on $N$ from Eq. 7 of the main text, we cancel the $v$ term and compare $\tau(\beta = 0)/\tau$. The expected increase in acquisition rate to keep the same phage MOI can then be found using Eq.25:

$$\begin{aligned} \frac{\tau(\beta=0)}{\tau^(\beta)} &= \frac{M\, N_b/\alpha N}{M\, N_b/\alpha N_\beta} \\ \frac{N_\beta}{N} &= \frac{r(\frac{1}{c^*}-1)}{r(\frac{1}{c^*_\beta}-1)} = \frac{1-c^*}{1-c^*_\beta} \times \frac{c^*_\beta}{c^*} \\ &= \frac{(1-c^*)e^{\frac{\beta\sigma^2}{r}}}{1-c^* e^{\frac{\beta\sigma^2}{r}}} \end{aligned} \tag{27}$$

Since $Q$ is a small correction and we only care about the scaling behavior, we can approximate $c^* \approx c^*_0 = 1 - R_0^{-1/M}$. To simplify the next derivation let $x = \beta\sigma^2/r$ and $q = 1 - c^* \approx R^{-1/M}$.

$$\begin{aligned} \frac{N_\beta}{N} &= \frac{(1-c^*)e^{\frac{\beta\sigma^2}{r}}}{1-c^* e^{\frac{\beta\sigma^2}{r}}} \\ &= \frac{qe^x}{1-(1-q)e^x} \\ \left(\frac{N_\beta}{N}\right)^{-1} &= \frac{1}{q}(e^{-x} - 1 - q) \\ 1 - \left(\frac{N_\beta}{N}\right)^{-1} &= \frac{1}{q}(1 - e^{-x}) \end{aligned} \tag{28}$$

Another simplification we should use is $z = \frac{1}{q}(1 - e^{-x})$. This is just to simplify when taking log of both sides:

$$\begin{aligned} \left(\frac{N_\beta}{N}\right) &= \frac{1}{1-z} \\ \ln\left(\frac{N_\beta}{N}\right) &= -\ln(1-z) \end{aligned} \tag{29}$$

We can then take the log of both sides of the equation and use $-\ln(1-z) = z + \mathcal{O}(z^2)$ and $1 - e^{-x} = x + \mathcal{O}(x^2)$ since $x = \beta\sigma^2/r$ is small (around $10^{-2} \sim 1$):

$$\begin{aligned} \ln\left(\frac{N_\beta}{N}\right) &= -\ln(1-z) \\ &\approx z = \frac{1-e^{-x}}{q} \approx \frac{x}{q} \\ &= R_0^{1/M}\beta\sigma^2/r \end{aligned} \tag{30}$$

### 5. Memory Fluctuation

Horizontal Gene Acquisition rate is modeled after a Poisson process. The expected time between HGT events is $T_{\mathrm{HGT}}$, and the memory size right after an HGT event is given by an exponential decay function. In this section, $N_0$ is specifically used to refer to the equilibrium phage population (in the absence of any perturbations.

$$M(t) = \gamma \frac{N_0}{N_b} e^{\frac{-t}{\tau_m}} + M_0 \quad [31]$$

**A. Initial Perturbation.** Under these constraints, the total number of spacers in the host (bacteria) population is given by the total time difference between the total number of spacers acquired $N^+$ and the total number of spacers lost $N^-$.

$$N_h * M(t) = \int N^+(t)\, dt - \int N^-(t)\, dt \quad [32]$$

Before we tackle the effect of differentiating acquisition and loss tradeoffs, we can first solve for the effect of a one-time increase in memory size and its effect on the overall fitness. Here, we assume a linear perturbation on the coverage immediately after the acquisition of $\gamma N/N_b$ ne memories. Here, we define $c_\delta$ as the coverage immediately after the increase in memory, which we take to be at $t = 0$ without loss of generality.

$$\int f(c)\, dt = \int f(c^*) + \partial_c f(c^*) |c_\delta - c^*|\, dt \quad [33]$$

Since $c^*$ is by definition the zero of the fitness function, we are only concerned with the second term of eq. 33. Since the perturbation on $n_s$ is just a net increase of $\gamma N$ over the baseline, the added perturbation due to the increase is an addition of $\gamma N e^{-t/\tau}\, \delta(x + vt)$ where we assume the perturbation occurs at $(x, t) = (0, 0)$ and $v$ is not significantly different from the unperturbed case. Evidence that $v$ is mostly unchanged can be seen in Fig.S7B. We can calculate $|c_\delta - c^*|$ through another ballistic approximation:

$$\begin{aligned} c_\delta &= \frac{1}{MN_b} \int n_s(x', t) e^{-|x'' - x'|/r}\, dx' \\ &= \frac{1}{MN_b} \int \left[ n_s(x', t; \gamma = 0) + \gamma N \delta(x' + vt) \right] e^{-|x'' - x'|/r}\, dx' \\ &= c^*(x, t; \gamma = 0) + \gamma \frac{N}{MN_b} e^{-|x'' + vt|/r} \\ &= c^* + \frac{\gamma \epsilon(x'')}{\alpha \tau} e^{-|vt|/r} \end{aligned} \quad [34]$$

This gives us the effect of the perturbation of the increased memory size. $\epsilon(x'')$ is the correction term because the surviving population will rarely feel the full effect of the added memories $\gamma N/MN_b$ since phages at $x'' \leq 0$ tend to immediately die and thus the rate of survival and fitness tend to be defined by the furthest phages away. In this case, we need to move the ballistic-like approximation away from the wave center by a distance of $x''$. In the plot of Fig.3B of the main text, we found the correction to be $\epsilon = 0.14$ to correct for such effects. We did not consider perturbations in $v$ and the rate of memory loss since we are only concerned with the short-term effects of this perturbation. In the previous Fig. 3B, the transient population size was found by averaging both the numeric and simulated phage population for 10 timesteps after the initial perturbation.

Finally, this leads us to two types of models for HGT: the acquisition-suppressed HGT, where $N^+$ is immediately suppressed after an HGT event, and the loss-enhanced HGT, where $N^-$ is enhanced immediately after an HGT event. In both cases, the conjugate loss/acquisition rate is taken as constant.

**B. Long-Term Calculations.** We want to find the long-term effect of losing more spacers per generation compared to the base case. From Eq.1, it follows that the solution for $n_s(x, t) = \alpha \int K(t, t')\, n(x, t')\, dt'$ where $K(t, t')$ is the memory kernel which represents the probability of retaining a spacer acquired at time $t'$ over time $t$. In the base case, the memory kernel takes the form of a simple exponential decay:

$$K(x, t) = \exp(-\frac{t - t'}{\tau}) \quad [35]$$

Solving for the long-term case, we are solving for the short-term recovery from the equation:

$$\partial_t n_s(x, t) = N^+(t) \frac{n(x, t)}{N} - N^-(t) \frac{n_s(x, t)}{M(t)\, N_b} \quad [36]$$

***B.1. Loss Enhanced.*** The loss-enhanced model of recovery is driven by a higher loss rate for a recovery timescale $\tau_m$ while keeping the acquisition rate constant:

$$
\begin{aligned}
N^+(t) &= \alpha N_0 \\
N^-(t) &= \alpha N_0(1 + \frac{\gamma}{\tau_m} e^{\frac{-t}{\tau_m}})
\end{aligned}
\tag{37}
$$

The solution for this equation is given by the memory kernel:

$$
\begin{aligned}
h^-(t) &= e^t \left(\gamma N_0 + M_0 N_b \exp(t/\tau_m)\right)^{\tau_m(1/\tau - 1)} \\
K^-(t, t') &= \frac{h^-(t')}{h^-(t)}
\end{aligned}
\tag{38}
$$

We can approximate $h^-(t)$ given that $\gamma/\tau \ll 1$ and $t/\tau_m \ll 1$ by:

$$
h^-(t) \approx \exp\left(-(\frac{1}{\tau} - 1)(1 - \frac{\gamma}{\tau}) + 1)t\right) \tag{39}
$$

In this loss-enhanced model, the approximate decay rate is increased compared to the base case, leading to a higher loss for a short amount of time after the fluctuation.

***B.2. Acquisition Suppressed.*** Conversely, the acquisition suppressed model is driven by lower acquisition over a similar recovery timescale $\tau$:

$$
\begin{aligned}
N^+(t) &= N_0(1 - \frac{\gamma}{\tau_m} e^{\frac{-t}{\tau_m}}) \\
N^-(t) &= N_0
\end{aligned}
\tag{40}
$$

Similarly, we have another memory kernel for this model:

$$
\begin{aligned}
h^+(t) &= \left(\gamma N_0(1 + \frac{\tau}{\gamma} \exp(t/\tau_m))\right)^{\frac{\tau_m}{\tau}} \\
K^+(t, t') &= \frac{h^+(t')}{h^+(t)} \left(1 - \frac{\gamma}{\tau_m} \exp(-t'/\tau_m)\right)
\end{aligned}
\tag{41}
$$

We can approximate $h^+(t)$ given that $\gamma/\tau \ll 1$ and $t/\tau_m \ll 1$ by:

$$
h^+(t) \approx \exp\left((-\frac{\gamma}{\tau^2} + \frac{1}{\tau})t\right) \tag{42}
$$

Compared to the base case, in the short term after the start of the fluctuation, the decay rate is comparatively smaller, but the probability of acquisition at $t = t'$ is not equal to 1 compared to the loss-enhanced and base cases.

**C. Damped Oscillatory Response to Fluctuations.** One very noticeable phenomenon when looking at the immediate response of the phage population after the fluctuation is a damped oscillatory response to the initial decay in fluctuation (Fig. S7). A damped response is a natural result of the higher-order dynamics of this system:

$$
\partial_t n(x, t) = D\partial_x^2 n(x, t) + f(x, t)n(x, t) \tag{43}
$$

Using the linear approximation of fitness from Ref. (1, 11), we can expand the fitness function in the traveling frame of reference $u$. Here, we left $f_0(t)$ as a time-dependent function to account for fitness effects not captured in the traveling frame of reference.

$$
\partial_t n(u) = D\partial_u^2 n(u) + (f_0(t) + su + \mathcal{O}(u^2))n(u) \tag{44}
$$

Taking the integral over $du$ and taking the time derivative again, the diffusion term goes away due to conservation of mass, and using the definition of $u = x - vt$, we obtain an expression for the harmonic motion of $N(t)$:

$$
\partial_t^2 N(t) = -svN(t) + \left(f_0(t) + \mathcal{O}(u^2)\right) \partial_t N(t) \tag{45}
$$

In the stable coevolutionary case, only the first term remains, as $f_0(t)$ and $\partial_t N$ are relatively small outside of some demographic noise. Under fluctuation, those terms are not zero, and thus we obtain a regime similar to a simple damped harmonic oscillator. However, these simple approximations have not been able to correctly predict the simulated period and amplitudes of driven oscillations in $N(t)$ but give us a good qualitative understanding of the dynamics at equilibrium. The expected period of this formulation is $T = 2\pi/\sqrt{sv}$, which is shorter than what we observe in simulation.

To get a better approximation of the time period of oscillations after a fluctuation, we should look at the interplay between coverage and phage population at the tip of the traveling wave: since the acquisition of new spacers is determined by the local abundance of phages, the spacer population also undergoes periods of low and high acquisition due to the periodic nature of the phages. When the spacer abundance is low because of the fluctuation, the next few timesteps will also have low acquisition, thus leading to a future period of high abundance. This phenomenon is similar to a classical predator-prey model, where we have periodic solutions. However, because of the dynamics of our system, the following systems of differential equations include a time delay in the coverage:

$$\begin{aligned} \frac{d\tilde{n}}{dt} &= (\underbrace{su_c}_{\text{fitness at tip of the wave}} - \underbrace{|\partial_c f(c^*)|\tilde{c}}_{\text{change in fitness from coverage}})\,\tilde{n} \\ \frac{d\tilde{c}}{dt} &= \underbrace{-\frac{v}{r}\tilde{c}}_{\text{decay in coverage due to wave travel}} + \underbrace{\frac{\alpha}{M\,N_b}\tilde{n}(t-u_c/v)}_{\text{spacer acquisition}} + \underbrace{\frac{\gamma}{\alpha\tau}e^{-|vt|/r}}_{\text{fluctuation recovery}} \end{aligned} \tag{46}$$

In Eq. 46, we have constructed a predator-prey-like model using $\tilde{n}$ and $\tilde{c}$, the phage population and immune coverage specifically at the tails of the traveling wave. All term found here arises naturally from previous theory. However, we have a time delay instead of a $\tilde{c}\tilde{n}$ cross-term in the dynamics to close the feedback loop. This system predicts an oscillatory period of $2\pi/\sqrt{svu_c/r}$, which is much closer than the previous approximation.

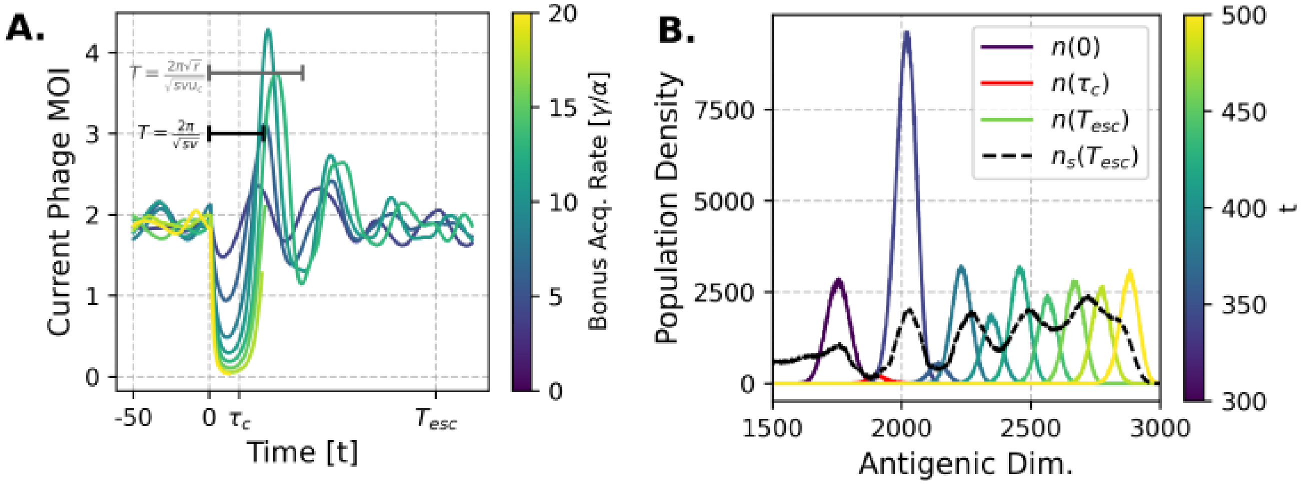


**Fig. S7.** Effects of Memory fluctuation. Simulation shown here was done with $N_b = 10^5$, $\mu = 10$, $M_0 = 5$. (A) Phage MOI overtime after an event at $t = 0$, the system oscillates back to equilibrium until around $T_{\text{esc}} = r/v$. Each line represents the average population over 20 runs, but most simulations at $\gamma/\alpha = 20$ will observe all phages going extinct. (B) Movement of the wave over time for $\gamma/\alpha = 10$, the pacer wave $n_s$ remembers the past motion of the phage wave and clearly illustrates the oscillatory pattern back to equilibrium. All phage waves, including $t = 0, \tau_c, T_{esc}$, were plotted with a 20 timesteps difference between them, showing us that $v$ is mostly constant, even as the population size fluctuates.
